# ATGs ubiquitination is required for circumsporozoite protein to subvert host innate immunity against malaria liver stage

**DOI:** 10.1101/2021.01.20.427456

**Authors:** Hong Zheng, Xiao Lu, Kai Li, Feng Zhu, Chenhao Zhao, Taiping Liu, Yan Ding, Yong Fu, Kun Zhang, Taoli Zhou, Jigang Dai, Yuzhang Wu, Wenyue Xu

**Affiliations:** Department of Pathogenic Biology, Army Medical University (Third Military Medical University), Chongqing 400038, China; The Institute of Immunology, Army Medical University (Third Military Medical University), Chongqing 400038, China; Department of Thoracic Surgery, Xinqiao Hospital, Army Medical University (Third Military Medical University), Chongqing 400037, China; Key Laboratory of Extreme Environmental Medicine, Ministry of Education of China, Chongqing 400037, China

**Keywords:** Plasmodium, malaria, sporozoites, exo-erythrocytic forms, circumsporozoite protein, IFN-γ, autophagy, ATGs, ubiquitination, E3 ligase

## Abstract

Although exoerythrocytic forms (EEFs) of liver stage malaria parasite in parasitophorous vacuole (PV) encountered with robust host innate immunity, EEFs can still survive and successfully complete infection of hepatocytes, and the underlying mechanism is largely unknown. Here, we showed that sporozoite circumsporozoite protein (CSP) translocated from the parasitophorous vacuole into the hepatocyte cytoplasm significantly inhibited the killing of exo-erythrocytic forms (EEFs) by interferon-gamma (IFN-γ). Attenuation of IFN-γ-mediated killing of EEFs by CSP was dependent on its ability to reduce the levels of autophagy-related genes (ATGs) in hepatocytes. The ATGs downregulation occurred through its enhanced ubiquitination mediated by E3 ligase NEDD4, an enzyme that was upregulated by CSP when it translocated from the cytoplasm into the nucleus of hepatocytes via its nuclear localization signal (NLS) domain. Thus, we have revealed an unrecognized role of CSP in subverting host innate immunity and shed new light for a prophylaxis strategy against liver-stage infection.

## Introduction

Malaria is still one of the most devastating diseases worldwide, which is caused by infection of the genus *Plasmodium*. Infection with *Plasmodium* is initiated in the mammalian host with the inoculation of sporozoites into the skin by *Anopheles* mosquitoes. From the skin, sporozoites reach the circulation and travel to the liver where they infect hepatocytes.

In the liver, sporozoites bind to highly-sulfated heparin sulfate proteoglycans (HSPGs) of hepatocytes surface by circumsporozoite protein (CSP) and trigger their invasion into hepatocytes (Coppi et al., 2007). After invading hepatocytes, sporozoites inside a parasitophorous vacuole (PV) transform into exo-erythrocytic forms (EEFs). To avoid the fusion of PV with lysosome and shield itself from the cytosol of the host, PV membrane is modified by several hepatocyte- and parasite-derived proteins (Kaushansky et al., 2015; Mueller et al., 2005; Sa et al., 2017; Silvie et al., 2003). However, EEFs in PV can be sensed by Melanoma differentiation-associated gene 5 (MD-5) and elicit hepatocytes to generate a type-I IFN response (Liehl et al., 2014), which recruits the innate immune cells, such as natural killer (NK) and NK T cells, and eliminates the EEFs in the infected hepatocytes through the secretion of IFN-γ (Miller et al., 2014). In addition, the infection of sporozoites could also induce the PAAR (Plasmodium associated autophagy-related) responses of hepatocytes to limit EEFs development(Akbari et al., 2018; Prado et al., 2015). However, EEFs can still survive and successfully complete the infection of hepatocytes, and the underlying mechanism of parasite survival in this hostile environment remains largely unknown.

Here, we found that CSP, the major surface protein of the sporozoites, translocating from the PV into the cytoplasm of hepatocytes (Singh et al., 2007), could resist the killing effect of IFN-γ on EEFs, and facilitate EEFs survival in hepatocytes. Therefore, we uncovered a novel immune escaping strategy of EEFs. Our further study showed that the resistance of CSP to the IFN-γ-killing of EEFs was dependent on its ability to downregulate autophagy-related genes (ATGs) through the enhanced ubiquitination mediated by E3 ligase NEDD4.

## Results

### CSP translocated into cytoplasm of hepatocyte remarkably resists the IFN-γ-mediated killing of EEFs

The pexel I-II domain of CSP has been demonstrated to mediate its translocation from the PV into the hepatocyte cytoplasm (Singh et al., 2007). To investigate whether CSP could suppress host innate immunity, a CSP pexel I-II domain mutant *Plasmodium berghei* (*P.b*) ANKA, named after CSP_mut_, was constructed by replacing the wild-type (WT) *CSP* with a pexel I-II mutant *CSP* using CRISPR-Cas9 technology (Figure S1A-C). As no mutations were found at any of the potential off-target regions screened by genome-wide comparison (Tab. S1), the potential of genomic off-target mutation generated by our CRISPR-Cas9 strategy could be excluded. The growth of the blood-stage mutant parasite was normal as compared to that of the WT parasite (CSP_wt_), and the salivary gland sporozoite of CSP_mut_ could be successfully generated when the mutant parasite infected mosquitoes (Figure S1D). Meanwhile, no difference of invasion ability into hepatocytes was found between CSP_mut_ and CSP_wt_ parasites (4.3±0.2% vs 4.0±0.1%) at 6 h post infection *in vitro*. Immunofluorescence assay with anti- upregulated in infective sporozoites gene 4 (UIS4), a PVM specific marker, and anti-CSP demonstrated that the CSP of mutant sporozoites was greatly inhibited to translocate from the PV into the hepatocyte cytoplasm when infecting HepG2 cells (Figure 1A).

**Figure 1.**
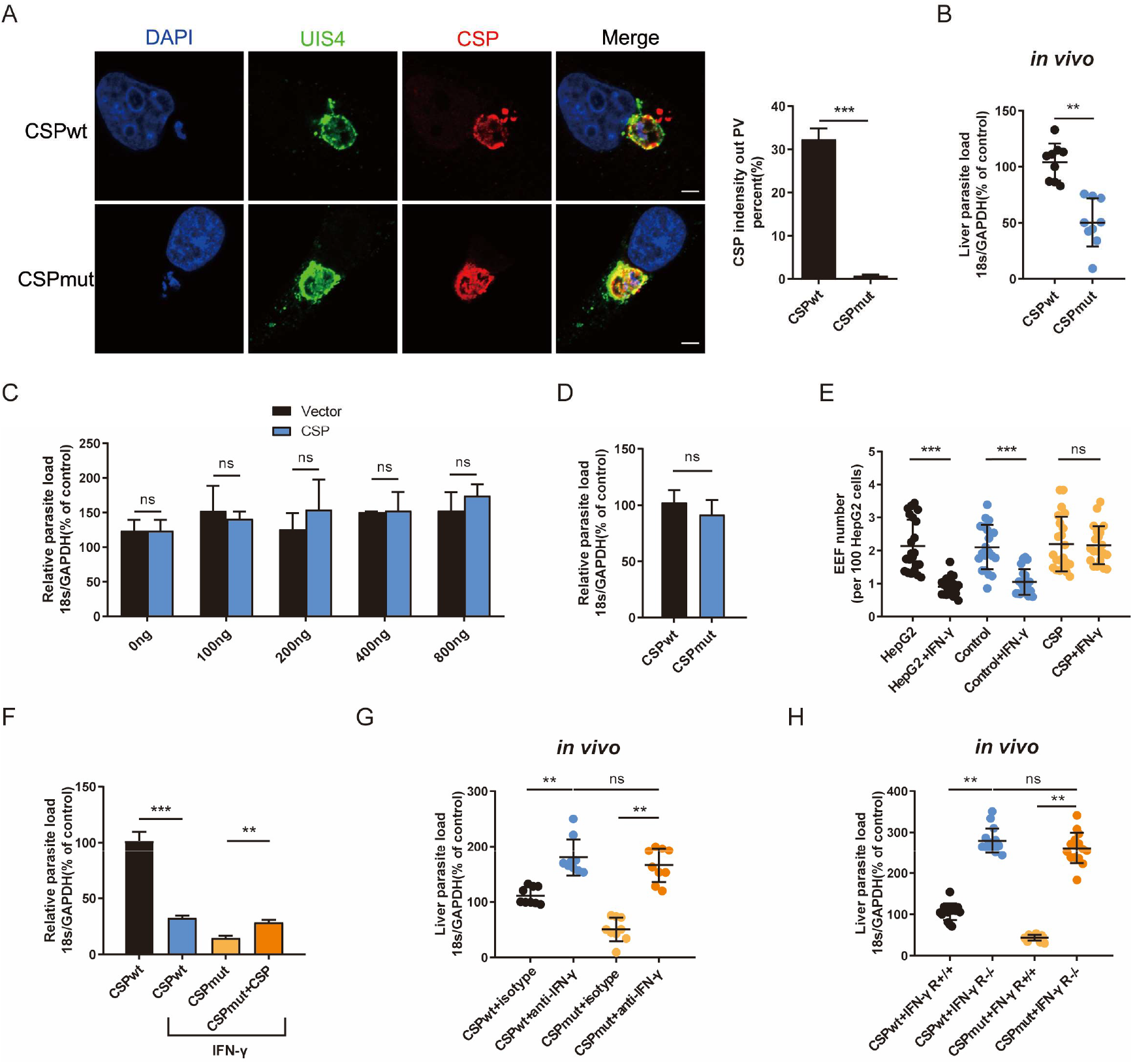
CSP translocated into cytoplasm of hepatocyte remarkably resists the killing of EEFs by IFN-γ. (**A**) After 1.2 × 10^5^ HepG2 cells were incubated with 4 × 10^4^ CSP_wt_ and CSP_mut_ *P.b* ANKA sporozoites for 24 h, then cells were stained with anti-UIS4, and anti-CSP and DAPI. Translocation of CSP from the PV into HepG2 cytoplasm was observed under confocal microscopy (*left*), and quantified (*right*). (**B**) After mice (n=5) were intravenously injected with 1,000 CSP_wt_ and CSP_mut_ parasite sporozoites for 46 h, the parasite burden in the liver was measured as the ratio of *Plasmodium* 18S rRNA to mouse *GAPDH* using Taqman real-time PCR. (**C**) HepG2 cells were transiently transfected with the indicated amount of pcDNA3.1 vector (Vec), CSP plasmid, and then incubated with sporozoites for 46 h; the parasite burden was determined as the ratio of *Plasmodium* 18S rRNA to human *GAPDH* using Taqman real-time PCR. (**D**) After HepG2 cells were incubated with CSP_wt_ and CSP_mut_ sporozoites for 46 h, the parasite burden was determined as described in (**C**). (**E**) Control and CSP-stable transfected HepG2 cells were pre-treated with or without 1U/ml IFN-γ, and then incubated with sporozoites for 46 h. EEF number was determined and statistically analyzed. (**F**) HepG2 cells were transfected with or without ΔCSP plasmid and pre-treated with or without 1 U/mL IFN-γ followed by incubation with WT or CSP_mut_ sporozoites; 46 h later, the parasite burden in hepatocytes was determined and compared as described in (**C**). (**G**) Mice (n=5) were infected with CSP_wt_ or CSP_mut_ sporozoites as above, and treated with anti-IFN-γ or isotype antibody. Then the parasite burden was determined as described in (**B**). (**H**) IFN-γR1 knockout and WT mice were infected with 1,000 CSP_wt_ or CSP_mut_ sporozoites, liver burden was determined at 46 h after infection as described as above. Med, Medium; Data are represented as mean ± SEM; ns, not significant; \*\**p* < 0.01; \*\*\**p* < 0.001. Scale bar=5μm.

Consistent with a previous study (Singh et al., 2007), we also found that the burden of CSP_mut_ in the liver was greatly reduced as compared to that of the CSP_wt_ after infected the mice (Figure 1B). However, transiently transfection with full-length CSP plasmid, mimicking CSP translocated into the cytoplasm of hepatocyte (Figure S2A-C), could not enhance the development of EEFs *in vitro*, even when up to 800 ng of plasmids were transfected (Figure 1C). There was also no significant difference in parasite load in HepG2 cells infected with CSP_wt_ or CSP_mut_ parasite *in vitro* (Figure 1D). As the infection of sporozoite could trigger mice to generate immune responses against parasite (Liehl et al., 2014), the different result between our *in vivo* and *in vitro* assay could be interpreted by the immune response induced *in vivo* that was not considered in our *in vitro* assay. Thus, we postulated that CSP might indirectly promote EEFs development through suppressing host immune responses.

*Plasmodium* in the liver could trigger a type I interferon (IFN) response and mainly recruits CD1d-restricted NKT cells (Miller et al., 2014). IFN-γ, predominately secreted by the recruited NKT cells, is considered as not only the major innate immune effector to suppress the development of the liver stage *in vivo* (Liehl et al., 2015; Miller et al., 2014), but also the critical effector to kill the EEFs in PV of hepatocytes *in vitro* (Ferreira et al., 1986; Schofield et al., 1987). Altogether, these findings strongly suggest that IFN-γ should be included in our investigation of the effect of CSP on EEFs development both *in vitro* and *in vivo.* Strikingly, we found that transfection of CSP plasmid greatly inhibited the IFN-γ-mediated killing of EEFs development, as both EEF number and parasite load in CSP-stable transfected HepG2 cells were much higher than those in control cells, when treated with IFN-γ (Figure 1E and Figure S2D). As expected, although mutant parasite in HepG2 cells was susceptible to IFN-γ-mediated killing, the stable transfection of CSP remarkably increased its resistance to IFN-γ (Figure 1F). Subsequently, this finding was further confirmed in a physiologically relevant context, as no significant difference of the liver parasite burden was found between the CSP_wt_ and CSP_mut_ -infected mice when IFN-γ was depleted (Figure 1G). Similar result was obtained when two parasite strains infected IFN-γR1 knockout mice (Figure 1H). The reduced parasite burden in CSP_mut_- infected mouse liver was not due to the capacity of mutant parasites to induce host to produce a much higher level of IFN-γ, because a comparable mRNA level of IFN-γ was found in the liver of mouse when infected with WT and CSP_mut_ sporozoites at different doses (Figure S2E). Thus, our data demonstrated that translocated CSP from the PV into the hepatocyte cytoplasm confers substantial resistance against EEF killing by IFN-γ.

### Nitric oxide (NO) is not involved in the suppression of the IFN-γ-mediated suppressing of EEFs by CSP

Previous studies have shown that IFN-γ prevents *Plasmodium* liver-stage development by inducing the expression of inducible nitric oxide synthase (iNOS), an enzyme required for the production of NO(Klotz et al., 1995; Mellouk et al., 1991; Sedegah et al., 1994). Hence, we sought to investigate whether inhibition of the IFN-γ-mediated suppressing of EEFs by CSP was dependent on the downregulation of iNOS and NO. We found that the transfection with CSP couldn’t reduce the mRNA level of iNOS or the concentration of NO in the infected-HepG2 cells treated with IFN-γ (Figure S3A). Similarly, both the level of iNOS and NO were comparable between CSP_wt_ and CSP_mut_ parasite-infected HepG2 cells treated with IFN-γ (Figure S3B). In addition, the resistance of CSP to the IFN-γ-mediated suppressing of EEFs, and the production of NO in CSP-transfected HepG2 cells treated with IFN-γ, were not affected by treatment with two iNOS inhibitors, aminoguanidine (AG) and L-NAME, although they could greatly inhibit the production of NO in lipopolysaccharide (LPS)-stimulated macrophages (Figure S3C). Therefore, NO is not involved in the suppression of the IFN-γ-mediated suppressing of the EEFs by CSP.

### Autophagy and its related proteins are essential for the IFN-γ-mediated killing of EEFs

Evidence has shown that autophagy and autophagy-related genes (ATGs) are essential for the IFN-γ-mediated killing of *Toxoplasma gondii* in macrophages (Choi et al., 2014; Ling et al., 2006; Ohshima et al., 2014; Zhao et al., 2008), an apicomplexan parasite closely related to *Plasmodium*. Thus, we next sought to investigate whether autophagy is also involved in regulation of the IFN-γ-mediated killing of EEFs in HepG2 cells. Consistent with our previous study (Zhao et al., 2016), autophagy, either enhanced by rapamycin or inhibited by LY294002, had no effect on liver-stage development in HepG2 cells (Figure 2A and B). However, rapamycin-enhanced autophagy greatly reduced either EEF number, parasite load or EEF size, in HepG2 cells treated with 0.2 U/mL IFN-γ, although this treatment alone had no effect (Figure 2A and Figure S4A). We found that rapamycin could not further reduce the parasite number and load, as well as size, when treated with 0.5 U/mL or 1 U/mL IFN-γ (Figure 2A and Figure S4A), which is possibly because most of the EEFs were already killed by IFN-γ at these higher concentrations (Schofield et al., 1987). As expected, the autophagy inhibitor LY294002 could remarkably reverse the killing effect of 0.5 U/mL and 1 U/mL IFN-γ on EEFs (Figure 2B and Figure S4B). A similar result was also obtained using another autophagy inhibitor, wortmannin (Figure 2C). No toxicity effect on cells was found for all above three drugs and IFN-γ at the working concentration (Figure S4C). Furthermore, the knockdown of either ATG5 (key component of ATG12-ATG5 conjugate) or ATG7(E1-like enzyme), by specific small hairpin RNA (shRNA) could significantly inhibit the IFN-γ-mediated suppressing of EEFs (Figure 2D and E). In addition, the depletion of IFN-γ greatly increased the liver parasite load in control ATG5^fl/fl^ mice, but had no effect on parasite burden in ATG5^fl/fl^-*Alb*-*Cre* mice, of which the ATG5 in hepatocytes is specifically knocked out (Figure 2F). These results demonstrated that autophagy and ATGs are pivotal for the killing of EEFs in hepatocytes by IFN-γ.

**Figure 2.**
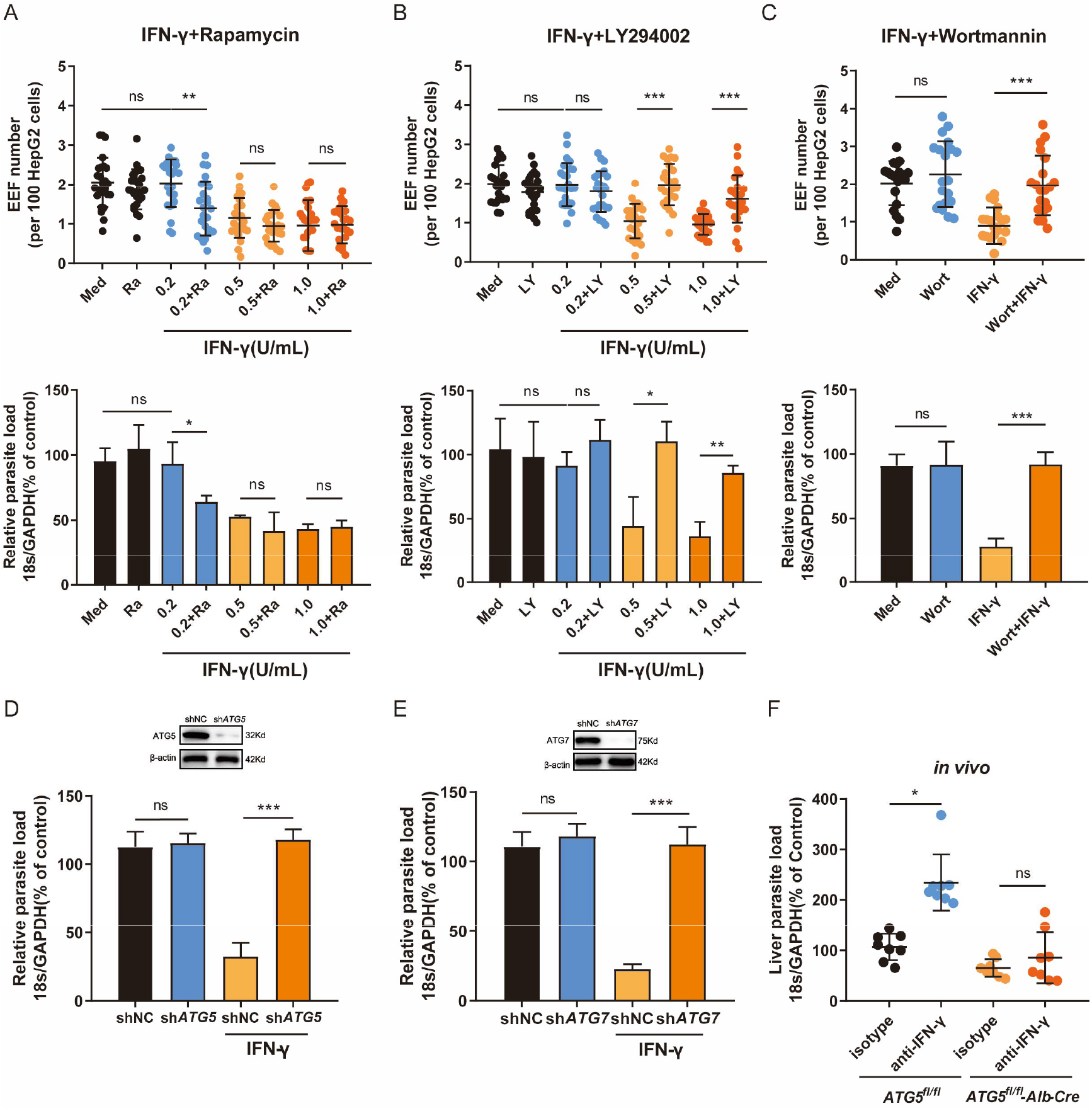
Autophagy and ATGs are essential for the IFN-γ-mediated killing of EEFs. (**A**) 1.2 × 10^5^ HepG2 cells were pre-treated with or without the autophagy inducer rapamycin (Rapa) and indicated concentrations of IFN-γ and then incubated with 4 × 10^4^ sporozoites. The EEF number (*up*) and the parasite load (*down*) at 46 h post-infection were compared. (**B**) HepG2 cells were pre-treated with or without the autophagy inhibitor LY294002 (LY) and IFN-γ at the indicated concentrations and then incubated with sporozoites. The EEF number (*up*) and the parasite load (*down*) at 46 h post-infection were determined. (**C**) HepG2 cells were pre-treated with or without the autophagy inhibitor wortmannin (Wort) and IFN-γ, and then incubated with sporozoites for 46 h. The EEF number (*up*) and the parasite burden (*bottom*) were determined. (**D**) HepG2 cells were transiently transfected with shNC or sh*ATG5* plasmids, and then treated with 1 U/mL IFN-γ and infected with sporozoites for 46 h. The knockdown of ATG5 was verified by western blot (*up*), and the parasite burden (*down*) in HepG2 cells was determined and compared. (**E**) HepG2 cells were transiently transfected with shNC or sh*ATG7* plasmids and then treated with IFN-γ and infected with sporozoites for 46 h. The knockdown of ATG7 was verified by western blot (*up*), and the parasite burden (*down*) in HepG2 cells was determined and compared. (**F**) After IFN-γ was depleted with or without anti-IFN-γ, both control (ATG5^fl/fl^) and ATG5^fl/fl^-*Alb*-*Cre* mice (n=5) were intravenously injected with 1,000 sporozoites, and then the liver parasite burdens were measured and compared. Data are represented as mean ± SEM; ns, not significant; \**p* < 0.05; \*\**p*<0.01; \*\*\**p*<0.001

### CSP greatly inhibits the IFN-γ-mediated suppression of EEFs by the downregulation of ATGs

Then, we investigated whether CSP could inhibit autophagy and the expression of ATGs. The rapamycin-induced autophagy was significantly inhibited in CSP-stable transfected HepG2 cells, since either the fluorescence intensity of key autophagy marker, microtubule-associated protein 1 light chain 3 (LC3)-RFP or LC3-GFP puncta, induced by rapamycin was reduced by more than 3 folds in CSP-stable transfected HepG2 cells as compared those of control cells (Figure 3A). In addition, the PAAR of EEFs in CSP-stable transfected HepG2 cells was also significantly inhibited as compared to that in the control cells (Figure 3B). In a physiologically relevant situation, we found that the intensity of LC3 fluorescence around the CSP_mut_ EEFs was much stronger than that of CSP_wt_ EEFs in HepG2 cells (Figure 3C). Next, the effect of CSP on the expression of ATGs was investigated. The LC3 level was greatly reduced in CSP-stable transfected HepG2 cells treated with rapamycin at all test time points, and the levels of autophagy-related genes, including Beclin-1, ATG3, ATG5, and ATG7, were also significantly decreased at 24 h. In contrast, the autophagy adaptor protein SQSTM1 (p62) significantly accumulated in CSP-stable transfected HepG2 cells, indicating the suppression of autophagy (Figure 3D). Overall, these findings demonstrated that CSP could inhibit autophagy and the expression of ATGs.

**Figure 3.**
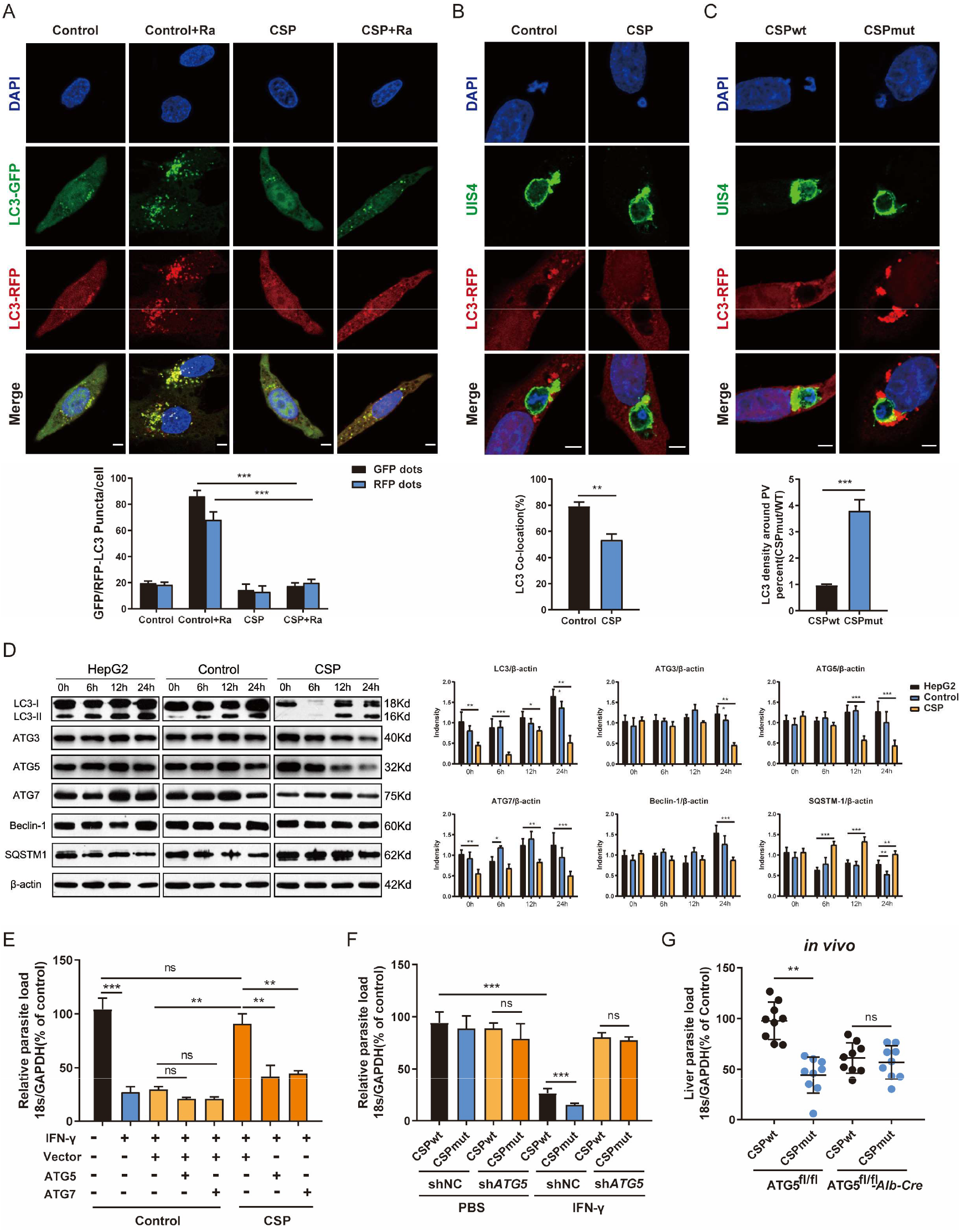
CSP greatly inhibits the IFN-γ-mediated suppression of EEFs by the downregulation of ATGs. (**A**) 1.2 × 10^5^ control and CSP-stable transfected HepG2 cells were infected with Ad-mRFP-GFP-LC3 virus, and then treated with the autophagy inducer rapamycin for 24 h. Both LC3-RFP or LC3-GFP puncta in HepG2 cells were imaged (*up*) and qualified (*down*). (**B**) Control and CSP-stable transfected HepG2 cells were transiently transfected with LC3-RFP plasmid, and then incubated with 4 × 10^4^ WT sporozoites for 24 h. Cells were fixed and stained with anti-UIS4, and LC3 surrounding the EEFs was observed under confocal microscopy (*up*), and the co-localization of LC3 and EEF was quantified (*down*). (**C**) HepG2 cells were transiently transfected with LC3-RFP plasmid and infected with CSP_wt_ or CSP_mut_ sporozoites for 24 h, The LC3 signals around WT and mutant EEFs were observed under confocal microscopy (*up*) and quantified (*down*). (**D**) Control and CSP-stable transfected HepG2 cells were treated with rapamycin for the indicated times. Protein levels of ATGs, including LC3I/LC3II, ATG3, ATG5, ATG7, Beclin-1, and p62 (SQSTM1), was determined by western blot (*left*), and the bands were quantified (*right*). (**E**) Control and CSP-stable transfected HepG2 cells transfected with or without ATG5, or ATG7 plasmid, and then treated with IFN-γ and infected with sporozoites for 46 h. The parasite load in HepG2 cells was determined and compared. (**F**) HepG2 cells were transiently transfected with Scramble or *ATG5* sh*RNA* plasmids, and then treated with IFN-γ and incubated with CSP_wt_ or CSP_mut_ sporozoites. 46 h later, the parasite load in the HepaG2 cells was determined and compared. (**G**) Both control (ATG5^fl/fl^) and ATG5^fl/fl^-*Alb*-*Cre* mice (n=5) were infected by intravenous injection with 1,000 WT or mutant sporozoites, 46 h later, the liver burden of the two parasites in control and ATG5^fl/fl^-*Alb*-*Cre* mice were determined and compared. Data are represented as mean ± SEM; ns, not significant; \*\**p* < 0.01; \*\*\**p* < 0.001. Scale bar=5μm.

As CSP could significantly suppress autophagy and its related proteins, we next investigated whether the resistance of CSP to IFN-γ-mediated suppression of EEFs was closely associated with its ability to suppress ATGs expression. Although the CSP-stable transfected HepG2 cells had a significant resistance to the IFN-γ-mediated suppression of EEFs, the overexpression of either ATG5 or ATG7 greatly abolished this inhibitory effect (Figure 3E). In addition, the burden of both CSP_wt_ and CSP_mut_ parasites in ATG5-knockdown HepG2 cells was comparable when the cells were treated with IFN-γ (Figure 3F). To further confirm this finding, both control (ATG5^fl/fl^) and ATG5^fl/fl^-*Alb*-*Cre* mice were infected with either CSP_wt_ or CSP_mut_ sporozoites, as expected, there was no difference in the liver burdens in ATG5^fl/fl^-*Alb*-*Cre* mice between two parasites (Figure 3G), indicating the essential role of ATGs in the resistance to the IFN-γ-mediated suppression by CSP *in vivo*. These findings demonstrated that ATGs are essential for the translocated CSP to resist the IFN-γ-mediated suppression of EEF development.

### The resistance of CSP to IFN-γ-mediated killing of EEFs is dependent on its ability to enhance ATGs ubiquitination

To explore the underlying mechanism by which CSP downregulates the expressing of the ATGs, mRNAs from CSP-stable transfected and control HepG2 cells treated with rapamycin were sequenced and compared. Unexpectedly, there was no significant down-regulation in mRNA levels of most ATGs in CSP-stable transfected HepG2 cells as compared to that of the control (Figure 4A), which was further confirmed by real-time PCR (Figure 4B). As CSP remarkably downregulated the protein levels of ATGs, this finding suggests that CSP does not regulate the expression of ATGs at the transcriptional level.

**Figure 4.**
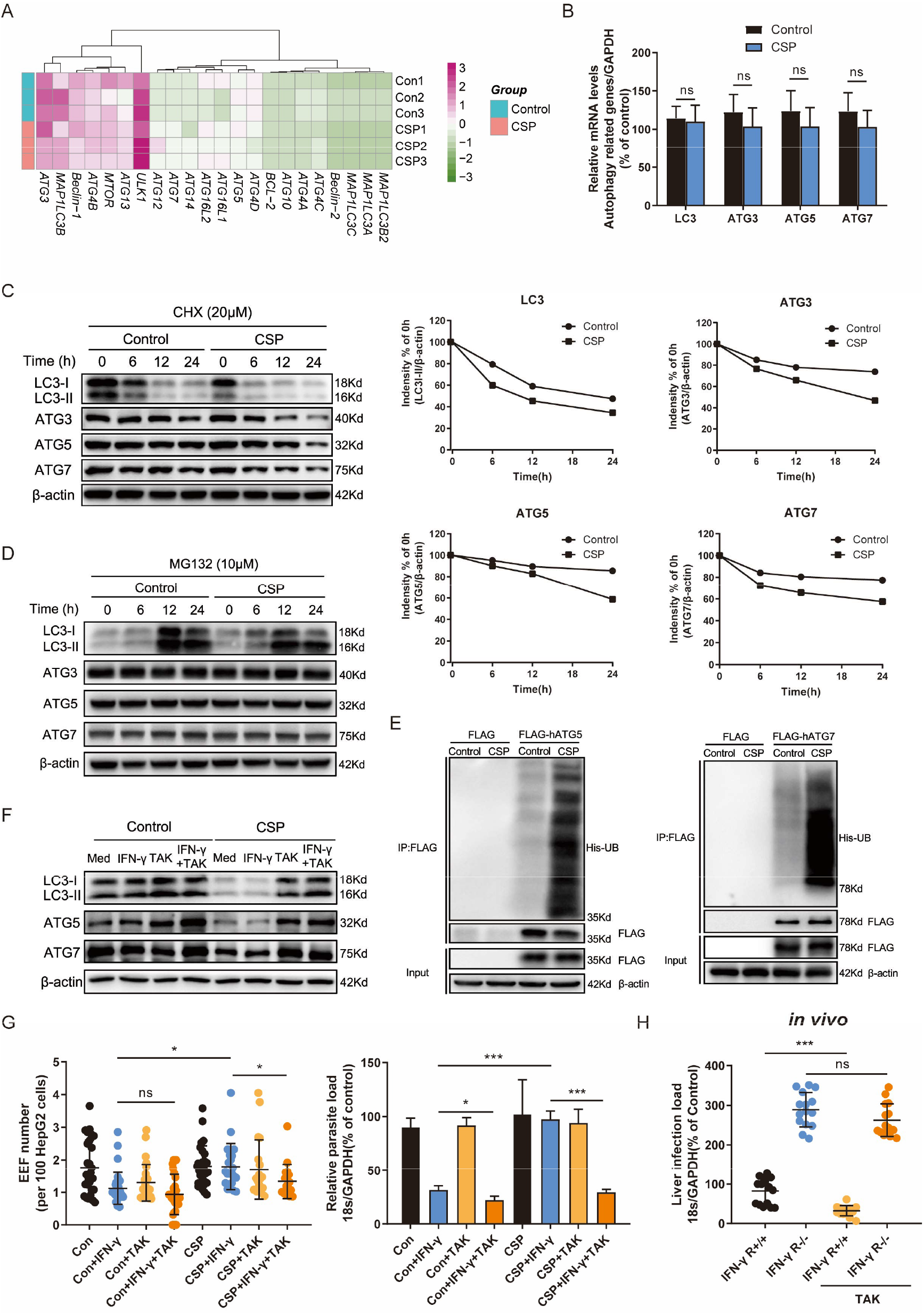
The resistance of CSP to IFN-γ-mediated killing of EEFs through the enhance of ATGs ubiquitination. (**A**) The mRNA levels of ATGs between control and CSP-stable transfected HepG2 cells in the heatmap were compared, and three biological repeats were performed. (**B**) The mRNA levels of ATGs (ATG3, ATG5, ATG7, and LC3) between control and CSP-stable transfected HepG2 cells were compared using real-time PCR. (**C**) Control and CSP-stable transfected HepG2 cells were treated with CHX for the indicated times, and the protein levels of LC3I/LC3II, ATG3, ATG5 and ATG7 were detected by western blot (*left*); the relative expression levels of LC3I/LC3II, ATG3, ATG5, and ATG7 to β-actin were quantified (*right*). (**D**) Control and CSP-stable transfected HepG2 cells were treated with the proteasome inhibitor MG132 for the indicated times, and the protein levels of LC3I/LC3II, ATG3, ATG5, and ATG7 were detected by western blot. (**E**) Control and CSP-stable transfected HepG2 cells were transfected with His-UB, and FLAG-ATG5 or FLAG-ATG7, and the ubiquitin binding to FLAG-ATG5 (*left*) or FLAG-ATG7 (*right*) in cells was detected by co-immunoprecipitation. UB, ubiquitin. (**F**) The levels of protein LC3, ATG5 and ATG7 were determined by western blot in both control and CSP-stable transfected HepG2 treated with or without 1μM TAK (TAK-243) for 24 h. (**G**) 1.2 × 10^5^CSP-stable transfected HepG2 or control cells were treated with or without 1μM TAK (TAK-243), and then were treated with IFN-γ and infected with 4 × 10^4^sporozoites for 46 h. The EEF number (*left*) and parasite load (*right*) were determined and compared as above. (H) IFN-γ R1 knockout and WT mice were per-treated with or without 20mg/kg TAK-243, and infected with 1,000 sporozoites in the next day, then liver parasite burden was determined at 46 h after infection as described as above. Data are represented as mean ± SEM; ns, not significant; \**p* < 0.05; \*\**p* < 0.01.

Eukaryotic cells use autophagy-lysosome and ubiquitin-proteasome system as their major intracellular protein degradation pathways(Varshavsky, 2017). However, autophagy level was outstandingly reduced in CSP-stable transfected HepG2 cells (Figure 3A), indicating that autophagy-lysosome system may be not responsible for the downregulation of the ATGs protein levels. Therefore, we next investigated whether CSP could downregulate the expression of ATGs at the post-translational level through ubiquitin-proteasome pathway, which is a major intracellular degradation pathway and a widespread post-translational modification(Hershko and Ciechanover, 1998). The half-lives of LC3, ATG3, ATG5, and ATG7 in the CSP-stable transfected HepG2 cells were significantly reduced compared to those of the control, when cells were treated with the protein synthesis inhibitor cycloheximide (CHX) (Figure 4C). In addition, the levels of LC3, ATG3, ATG5, and ATG7 in CSP-stable transfected and control HepG2 cells were comparable, when the cells were treated with the specific proteasome inhibitor MG132 (Figure 4D). These findings indicated that CSP could regulate the degradation of ATGs in a proteasome-dependent manner. To investigate whether this degradation of ATGs in the proteasome was dependent on ubiquitination, the CSP-stable transfected and control cells were transfected with His-UB, and FLAG-ATG5 or FLAG-ATG7 plasmids, and co-immunoprecipitation was performed using anti-FLAG beads to detect ubiquitin bound to ATGs. More ubiquitin were pulled by ATG5, especially ATG7, in the CSP-stable transfected HepG2 cells than that in control cells (Figure 4E). To verify the resistance of CSP to IFN-γ-mediated suppression of EEFs development was due to the enhancement of ATGs ubiquitination, a ubiquitination specific inhibitor TAK-243 (Hyer et al., 2018), was used at non-toxicity concentration (Figure S5) to suppress ubiquitination. We found that the treatment with TAK-243 significantly elevated the protein levels of ATGs (Figure 4F), and enhanced the IFN-γ-mediated killing of EEFs in CSP-stable transfected HepG2 cells, as compared to those in control cells (Figure 4G). Furthermore, the administration of TAK-243 could also greatly reduced the liver parasite burden in WT mice (n=5), but has no significant effect on liver parasite burden in IFN-γ R1 knockout mice (n=5) (Figure 4H). Thus, we demonstrated that CSP resists to IFN-γ-mediated killing of EEFs through the upregulation of specific ubiquitination of ATGs.

### CSP upregulates the E3 ubiquitin ligase NEDD4 to promote the ubiquitination of ATGs

E3 ubiquitin ligase, which transfers the ubiquitin of E2 enzyme to its attached substrate, is the key enzyme of the ubiquitination process (Hershko and Ciechanover, 1998). Therefore, we hypothesized that E3 ubiquitin ligases might be involved in the ubiquitination of ATGs (Li et al., 2017). Six candidate E3 ligases, including STUB1 (STIP1 homology and U box-containing protein 1), SYVN1 (Synovial apoptosis inhibitor 1), CBL (Cbl proto-oncogene), SMURF1 (SMAD specific E3 ubiquitin protein ligase 1), PAFAH1B1 (Platelet activating factor acetylhydrolase 1b regulatory subunit 1), and NEDD4 (neural precursor cell expressed, developmentally down-regulated 4), were predicted to regulate ubiquitination of at least two ATGs using an online prediction tool (Figure S6 and Tab. S2) (Li et al., 2017). However, the RNA-seq data showed that only NEDD4, among these ligases, was significantly upregulated in CSP-stable transfected HepG2 cells (Figure 5A), and the increase of both the mRNA and protein levels of NEDD4 was confirmed by real-time PCR and western blot, respectively (Figure 5B and C). Furthermore, infection with the CSP_wt_ parasite, but not the CSP_mut_ parasite, significantly elevated the mRNA level of *NEDD4* in HepG2 cells (Figure 5D), strongly suggesting that the translocation of CSP from the PV into the hepatocyte cytoplasm could upregulate the expression of E3 ubiquitin ligase NEDD4. Like our results, upregulation of NEDD4 in sporozoite infected hepatocyte cell lines agreed with other’s findings in *P. b* ANKA infection model (data from GSE78931 and GSE72049) (Figure 5E). Moreover, the knockdown of *NEDD4* by shRNA could significantly reduce the parasite load in CSP-stable transfected HepG2 cells treated with IFN-γ (Figure 5F), indicating that NEDD4 was essential for the resistance of CSP to IFN-γ-mediated suppressing of EEFs.

**Figure 5.**
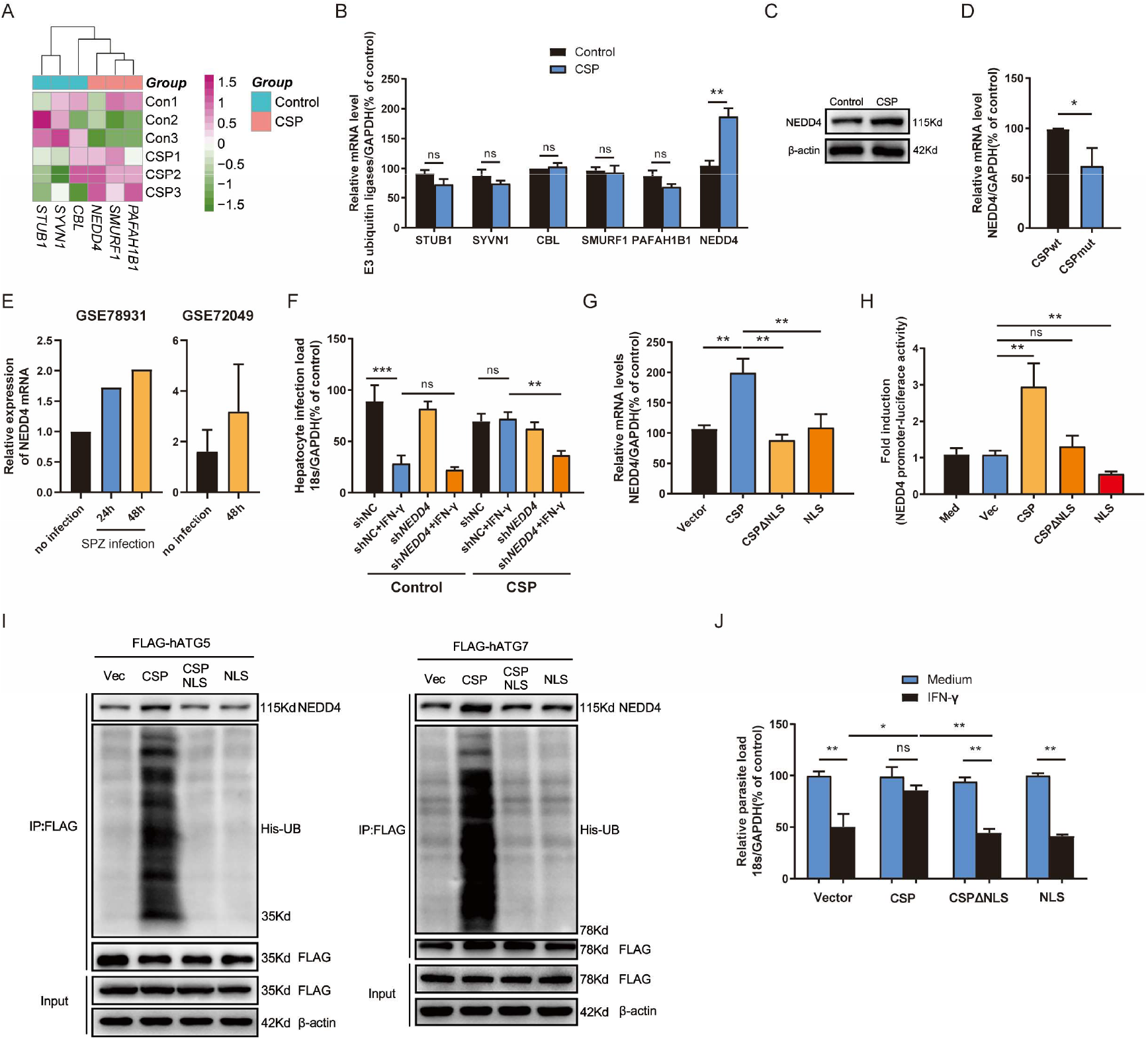
CSP upregulates the E3 ubiquitin ligase NEDD4 to promote the ubiquitination of ATGs. (**A**) The mRNA levels of six E3 ubiquitin ligases between control and CSP-stable transfected HepG2 cells in the heatmap were compared, and three biological replicates were performed. (**B**) The mRNA levels of six E3 ubiquitin ligases in both control and CSP-stable transfected HepG2 cells were confirmed by real-time PCR. (**C**) The protein levels of NEDD4 in both control and CSP-stable transfected HepG2 cells were confirmed by western blot. (**D**) After HepG2 cells were infected with CSP_wt_ or CSP_mut_ sporozoites for 24 h, the mRNA level of *NEDD4* was determined by real-time PCR. (**E**) Relative expression of NEDD4 mRNA in *P.b* ANKA sporozoite infected Huh7 and HepG2 cells were analyzed based on data extracted from GSE78931 and GSE 72049. (**F**) 1.2 × 10^5^ control and CSP-stable transfected HepG2 cells were transfected with shNC or sh*NEDD4* and treated with IFN-γ or not, then infected with 4 × 10^4^ sporozoites, for 46 h. The parasite load in HepG2 cells was determined as above. (**G**) HepG2 cells were transfected with CSP, CSPΔNLS, or NLS plasmids for 24 h, and the transcription of *NEDD4* was determined by real-time PCR. (**H**) HEK293T cells were transfected with RL-TK, *NEDD4* promoter reporter plasmid, and with or without CSP, CSPΔNLS, or NLS plasmids for 24 h, and the ratio of firefly luciferase to Renilla luciferase were determined using Dual Luciferase Assay kit. (**I**) 293FT cells were transiently transfected with His-UB, and vector, CSP, CSPΔNLS or NLS plasmids, and FLAG-ATG5 or FLAG-ATG7, and then treated with MG-132 for 24 h, both the NEDD4 and ubiquitin binding to FLAG-ATG5 (*left*) or FLAG-ATG7 (*right*) in cells were detected by co-immunoprecipitation. UB, ubiquitin. (**J**) 1.2 × 10^5^ HepG2 cells were transfected with vector, CSP, CSPΔNLS, or NLS plasmids and treated with or without IFN-γ, followed by the infection with 4 × 10^4^ sporozoites for 46 h. The parasite load in HepG2 cells was determined as above. Data are represented as mean ± SEM; ns, not significant; \**p* < 0.05; \*\**p* < 0.01.

A previous study showed that CSP contains a nuclear location signal (NLS) domain, which could import the CSP into the nucleus (Singh et al., 2007). To better elucidate the mechanism of the upregulation of NEDD4 by CSP, the effect of CSP, CSPΔNLS (CSP without NLS domain), and NLS (only NLS domain) (Figure S2A) on the transcription of *NEDD4* in HepG2 cells was determined. Although transfection of CSP could upregulate *NEDD4* expression, neither transfection of CSPΔNLS nor NLS influenced the transcription of *NEDD4* in HepG2 cells (Figure 5G). Dual luciferase reporter assay confirmed that only CSP could significantly activate the promoter of *NEDD4* (Figure 5H). These data indicated that the translocation into the nucleus is essential for CSP to modulate the transcription of *NEDD4*, but the modulatory motif might exist in the sequence of CSPΔNLS. In addition, co-immunoprecipitation showed that more NEDD4 proteins and ubiquitin were bound to both ATG5 and ATG7 in CSP, but not in CSPΔNLS or NLS-transfected 293FT cells (Figure 5I), and the transfection of CSP can only remarkably resist the IFN-γ-mediated suppression of EEFs in HepG2 cells (Figure 5J). Overall, these data suggested that NEDD4 is the E3 ubiquitin ligase to specifically mediate the ubiquitination of ATG5 and ATG7, and that CSP enhances the ubiquitination of ATGs through the upregulation of NEDD4.

## Discussion

Great progress has been made to identify the molecules of sporozoites involved in their selective attachment to hepatocytes and the formation of the PV after their invasion (Coppi et al., 2007; Kaushansky et al., 2015; Silvie et al., 2003; Yalaoui et al., 2008), however, the underlying mechanism of the survival of parasites in the PV has thus far remained unknown. Here, we found that CSP translocated from the PV to the cytoplasm could inhibit the IFN-γ-mediated killing of EEFs to facilitate the survival of parasites in the PV.

CSP is a multifunctional protein of the malaria parasite that is not only critical for the invasion of sporozoites into hepatocytes but is also essential for sporozoite development in mosquitoes (Cerami et al., 1992; Coppi et al., 2011; Frevert et al., 1993; Menard et al., 1997; Pinzon-Ortiz et al., 2001; Rathore et al., 2002; Wang et al., 2005). Therefore, CSP is regarded as a leading candidate protective antigen (Kumar et al., 2006), which is included in several subunit vaccines to induce CSP-specific antibody and/or CD8^+^T cells responses against sporozoites (Aliprandini et al., 2018; Foquet et al., 2014; Gordon et al., 1995; Kisalu et al., 2018; Kubler-Kielb et al., 2010; Stoute et al., 1997; Tan et al., 2018). Our findings of this new aspect of CSP to subvert the host innate immune response not only support the important role of CSP as the dominant protective antigen but further shed new light into a novel prophylaxis strategy against the liver stage.

Recent studies have demonstrated that infection of sporozoites could induce both the canonical autophagy and the PAAR response of hepatocytes against EEFs (Prado et al., 2015; Zhao et al., 2016). Although canonical autophagy promotes parasite development by supplying essential nutrients (Grutzke et al., 2014), the role of PAAR response in liver stage development is conflicting (Prado et al., 2015; Thieleke-Matos et al., 2016). One possible reason is the conditional ATGs knockout of non-permissive mouse embryonic fibroblasts (MEFs) was used in previous studies (Prado et al., 2015; Thieleke-Matos et al., 2016). In this study, we found the knockdown of either ATG5, or ATG7 in HepG2 cells, the permissive cells of sporozoites, has no significant effect on EEFs development (Figure 2D and E). However, the parasite load in ATG5^fl/fl^-*Alb*-*Cre* mice, of which ATG5 was completely knocked out in hepatocytes, was significantly lower than that of control mice, when IFN-γ was depleted (Figure 2F). This suggested that the fundamental level of autophagy was needed for EEF development. In addition, our findings that the induction of autophagy by rapamycin enhances the killing effect of IFN-γ (Figure 2A), and autophagy inhibitors or knockdown of ATGs attenuates the killing of EEFs by IFN-γ (Figure 2B and F), demonstrate the essential role of autophagy in the regulation of the IFN-γ-mediated killing of EEFs. This is supported by a recent finding that the downstream ATGs mediating LC3-associated phagocytosis (LAP)-like process regulate the killing of *Plasmodium vivax* liver stage by IFN-γ (Boonhok et al., 2016). Both IFN-γ-inducible GTPase Irg6 and guanylate-binding protein (GBP)1 have been identified to regulate the IFN-γ-mediated killing of intracellular *T. gondii* by recruitment to the PVM,(Liesenfeld et al., 2011; Selleck et al., 2013; Virreira Winter et al., 2011) but Irg6 is not involved in the IFN-γ-mediated killing of EEFs (Liesenfeld et al., 2011). Indeed, we found that the knockout of GBP-1 also had no effect on EEFs development in HepG2 cells treated with IFN-γ (Figure S7). Thus, the roles of GTPase and GBPs in the IFN-γ-mediated killing of EEFs might differ from those in *T. gondii,* and should be investigated in further detail.

Recent studies showed that parasites can shed host autophagic proteins from the parasitophorous vacuole membrane (PVM) to escape the deleterious effects of PAAR response (Agop-Nersesian et al., 2017) and *UIS3* sequesters the LC3 to suppress the autophagy of EEFs (Real et al., 2017). These findings did support the existence of subversive strategies of EEFs against hepatocyte autonomous immunity. However, we found that CSP translocated from the PV into the cytoplasm could inhibit the IFN-γ-mediated killing of EEFs through downregulating the expression of ATGs by ubiquitination, revealing a novel strategy of malaria liver stage to subvert host innate immunity.

Taken together, we found that CSP translocated from PV into cytoplasm of hepatocyte could be imported into nucleus and upregulates the transcription of E3 ligase NEDD4 dependent on its NLS domain. E3 ligase NEDD4 enhances the ubiquitination of ATGs, leading to the decrease of the protein level of ATGs in hepatocytes. As ATGs are essential for IFN-γ-mediated killing of EEFs, CSP resists to IFN-γ-mediated killing of EEFs, and facilitates the survival of EEFs in hepatocytes (Figure 6). Thus, our findings explained why CSP translocated from PV into the hepatocyte cytoplasm can promote the liver stage development, and provide novel prophylaxis strategies to eliminate liver stage infection through chemically targeting CSP pexel or NLS domain, or ubiquitination in hepatocytes.

**Figure 6.**
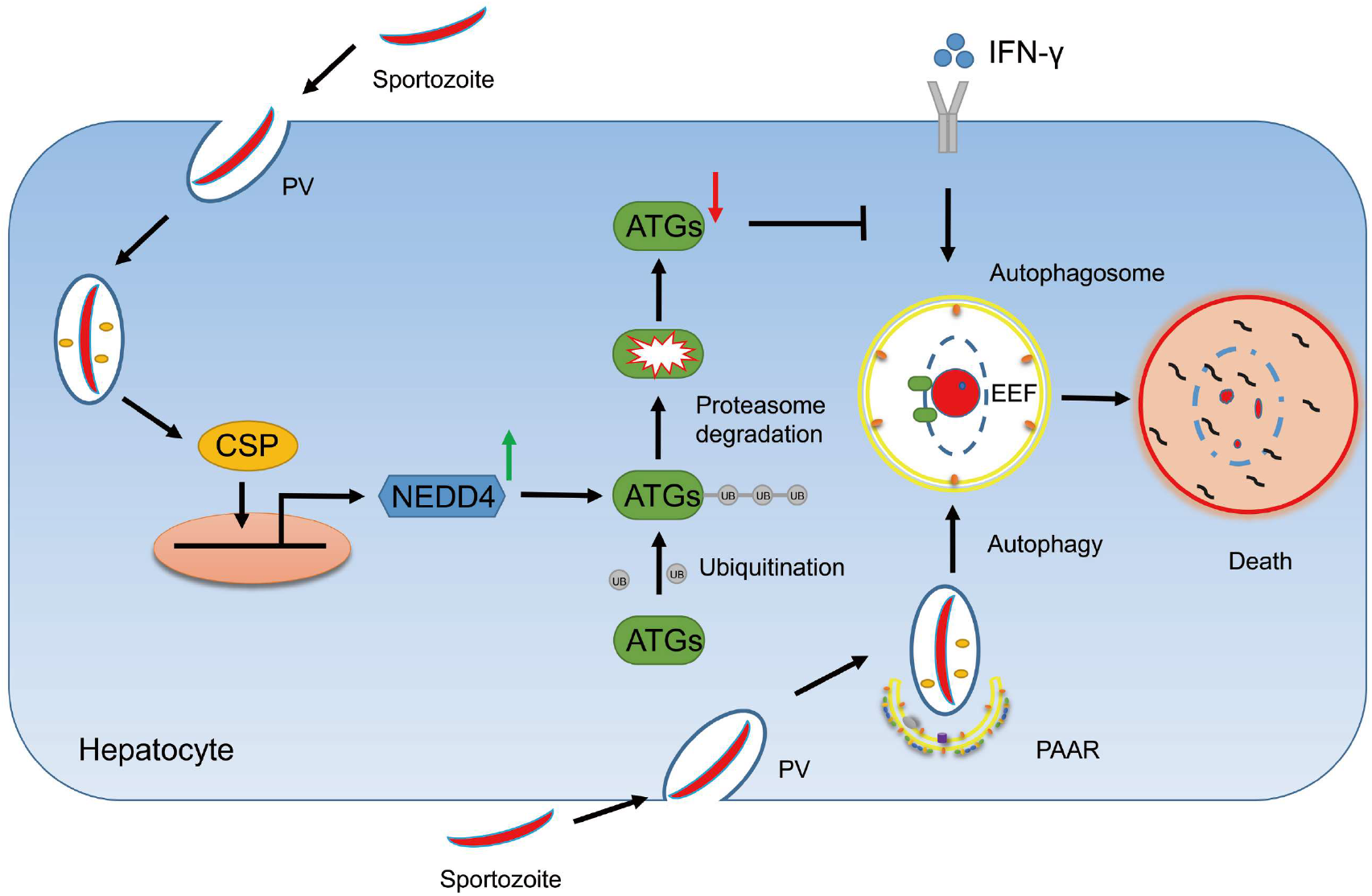
A mechanistic pathway of the resistance of CSP to IFN-γ-mediated killing of EEFs in hepatocytes. After sporozoite invaded into hepatocytes, CSP on its surface would be translocated from PV into the cytoplasm of hepatocytes, and is then introduced into nucleus dependent on its NLS domain. As a result, CSP activates the promoter of ligase3 *NEDD4* and enhances its transcription. The upregulated NEDD4 promotes the ubiquitination of ATGs, leading to the degradation of ATGs. As ATGs are essential for IFN-γ-mediated killing of EEFs, the resistance of CSP to IFN-γ, which is dependent on its ability to downregulate ATGs through enhancing NEDD4-mediated ubiquitination, facilitates the survival of EEFs. Otherwise, EEFs are destroyed by PAAR (Plasmodium associated autophagy-related).

## Material and Methods

### Parasites and mice

*Plasmodium berghei* ANKA (*P.b* ANKA) and *P.b* ANKA-RFP were maintained in our laboratory, and the *P.b* ANKA CSP_mut_ parasite was constructed by mutation of pexel I-II of CSP using CRISPR-Cas9. All parasites were maintained by passage between Kunming mouse (a Swiss Webster strain) and *Anopheles stephensi*. ATG5^fl/fl^ mice (B6.129S-Atg5<tm1Myok>) were obtained from RIKEN BioResource Center (Ibaraki, Japan), and Alb-Cre mice (B6.Cg-Speer6-ps1Tg(Alb-cre)21Mgn/J) were purchased from Jackson Laboratory (Bar Harbor, ME, USA). ATG5^fl/fl^ mice were crossed with *Alb-Cre* mice to conditionally delete the ATG5 cassette in hepatocytes. ATG5^fl/fl^ mice were used as control. IFN-γR1 knockout mice were gifts from Dr. Bo Guo (Army Medical University, Chongqing, China). Kunming mice were purchased from the Experimental Animal Center of the Army Medical University (Chongqing, China). Animals were kept in the specific pathogen-free laboratory of the Institute of Immunology of Army Medical University. All methods were carried out in accordance with the approved Guide for the Care and Use of Laboratory Animals of the Army Medical University. All experimental protocols were approved by the Animal Institute of Army Medical University.

### Mosquito rearing and infection

*Anopheles stephensi* (Hor strain) were maintained at 27°C, 70–80% relative humidity, and fed with a 5% sugar solution. For infection with the parasite, 3-to 5-day-old female adults were kept at 20–21°C, 70–80% relative humidity and fed on *P.b* ANKA, *P.b* ANKA-RFP, or *P.b* ANKA-CSP_mut_-infected Kunming mice with gametocytemia ≥ 0.5%. At 7 days following infection, the mosquitoes were dissected, and the oocysts on the midguts were examined under a light microscope. Sporozoites were isolated from the salivary gland at 19 days post-infection.

### Cell culture, transfection, and virus infection

Both HepG2 and HEK293T cells were purchased from American Type Culture Collection (Manassas, VA, USA). HepG2 cells were maintained in DMEM (Hyclone) and HEK293T cells were maintained in RPMI-1640 medium (Hyclone), and supplemented with 10% FBS and 1% penicillin/streptomycin (Gibco) at 37°C in a humidified atmosphere with 5% CO2. For transfection, 3 × 10^4^ cells in 96-well, 1.2 × 10^5^ cells in 24-well, 1 × 10^6^ cells in 6-well plates, and 1 × 10^7^ cells in 10-cm dishes were transfected with 0.2 μg, 0.8 μg, 5 μg or 24 μg recombinant plasmid using 2 μL, 7.5 μL or 37.5 μL Lipofectamine 3000 (Invitrogen, CA, USA), respectively, according to the manufacturer’s protocol. For virus infection, 1 × 10^5^ HepG2 cells in 24-well plates were pretreated with 8 μg/mL polybrene (Santa Cruz, CA, USA) for 2 h, and then infected with Ad-mRFP-GRP-LC3 (Hanbio, Shanghai, China) at a multiplicity of infection (MOI) of 15. Four hours later, fresh cell culture containing 8 μg/mL polybrene was added. After 12 h of virus infection, the supernatant was discarded and replaced with fresh culture medium.

### Infection of *P.berghei* ANKA sporozoites

At 19 days following infection with *P.b* ANKA, *P.b* ANKA-RFP, or *P.b* ANKA-CSP_mut_, infected female mosquitoes were extensively washed in sterile PBS, and the salivary glands were dissected and collected in RPMI 1640 medium containing 2.5 μg/mL amphotericin B (Sangon Biotech, Shanghai, China), 100 units/mL penicillin, and 100 μg/mL streptomycin (Beyotime, Beijing, China). The sporozoites were released from salivary glands and counted, then incubated with HepG2 cells at a ratio of 1:3. Three hours after infection, the supernatant was discarded and replaced with fresh culture medium with the three antibiotics as described above. For the *in vivo* assay, ATG5^fl/fl^ and ATG5^fl/fl^-*Alb-Cre* were challenged with intravenous injection of 1,000 sporozoites. Control and IFN-γ R1 knockout mice were challenged with intravenous injection of 10,000 sporozoites.

### Construction of recombinant plasmids

The fragments of *P.b* ANKA full CSP (1020bp) was amplified from the genome of *P.b* ANKA using KOD-FX DNA polymerase (TOYOBO, Osaka, Japan). CSPΔNLS (993 bp) and NLS (27 bp) fragments with initiation codons were synthesized by Sangon Biotech. CSP and CSPΔNLS with a 16-18 bp homologous sequence of the pcDNA3.1 vector at either the 5′ or 3′ end were obtained by PCR. The pcDNA3.1 vector was linearized using the restriction enzymes *Hind*III and *BamH*I. The fragments of CSP and CSPΔNLS were inserted into the pcDNA3.1 vector, respectively, by seamless cloning according to the instruction of In-fusion^®^ HD Cloning Kit (Clontech, Palo Alto, CA, USA). NLS was inserted into pcDNA3.1 using High-Efficient Ligation Reagent Ligation High (TOYOBO). pcDNA3.1 vector was linearized using the restriction enzymes *BamH*I and *Xhol*I (TAKARA, Otsu, Japan) for *ATG5 and LC3B*, and with *Kpn*I and *Not* I for *ATG7*. The pFLAG-cmv8 vector was linearized using the restriction enzymes *Hind*III and *EcoR*I for *ATG5* and *ATG7*. Both hLC3B, hATG5 and hATG7 were inserted into the pcDNA3.1 or pFLAG-cmv8 vector, respectively, by seamless cloning according to the instruction of In-fusion^®^HD Cloning Kit (Clontech). pGL3-NEDD4 promoter report plasmid was constructed by cloning NEDD4 promoter sequence (2000bp upstream form 1^st^ exon) into pGL3 vector though *Kpn*I/*Hind* III. All recombinant plasmids were verified by DNA sequencing.

### Construction of CSP-stable transfected cell lines

The CSP coding sequence of *P.b* ANKA was cloned downstream of the CMV-7 promoter in the Lenti-pCDH plasmid (a gift from Professor Ying Wan, Army Medical University, China). HEK293T cells were transfected with Lenti-pCDH-CSP or Lenti-pCDH control plasmids with psPAX2 and pMD2.G packaging plasmids as ratio 20:15:5 using Lipofectamine 3000 (Invitrogen) to obtain the lentivirus. Seventy-two hours after transfection, the supernatant was filtered with a 0.45-μm filter (Milipore) and the lentivirus was collected by ultracentrifugation (20,000 rpm) at 4°C for 2 h. HepG2 cells were infected with Lenti-pCDH-CSP or Lenti-pCDH control lentivirus at a MOI of 15 for 48 h. Then, the positively infected cells were screened by 5 mg/mL puromycin (Gibco) and sorted by flow cytometry. CSP-stable transfected and control cells were finally cloned by a limited-dilution method continually in a 96-well plate, and 3 mg/mL puromycin was used to maintain the resistance of stable transfected cells. Expression of CSP was confirmed by real-time PCR and immunofluorescence.

### Preparation of CSP antibodies

The peptide containing three *P.b* ANKA CSP repeats, PAPPNANA-PAPPNANA-PAPPNANA, was synthesized by Wuhan GeneCreate Biological Engineering Co., Ltd. (Wuhan, China), and purified by high-performance liquid chromatography (HPLC) using a Waters XBridge C18 column (4.6 × 250 mm × 5 μm; Waters Corporation) on a Waters Corporation Prominence HPLC system (Massachusetts, USA).

The peptide was conjugated with bovine serum albumin (BSA). The resulting conjugated peptide was emulsified with Freund's adjuvant (Sigma Aldrich) and immunized in a specific pathogen-free-grade rabbit at 0, 2, 4 and 6 weeks. Seven days after the final immunization, rabbit serum was collected, and the titer of the antibody against the epitope was detected using an enzyme-linked immunosorbent assay (cat. no., 44-2404-21, Nunc MaxiSorp flat bottom, Nalge Nunc International, Penfield, NY, USA). Protein G Sepharose (GE Healthcare) was used for purification of CSP antibody.

### Immunofluorescence assay

HepG2 cells (1 × 10^5^) were placed on a 14-mm-diameter slide in a 24-well plate. To detect the distribution of CSP, HepG2 cells were infected with CSP_wt_ or CSP_mut_ sporozoites. Three hours after invasion, the cells were washed with PBS to remove non-invaded sporozoites and incubated with fresh DMEM (Hyclone) containing 10% FBS (Gibco) and three antibiotics. Twenty-four hours after invasion, the slide was immobilized with 4% paraformaldehyde (Sangon Biotech) for 10 min, penetrated with 0.1% Triton-X 100 (Amersco, Albany, NY, USA) for 15 min, and then blocked with PBS containing 5% BSA, 0.02% Tween-20 for 1 h at room temperature. The cells were labeled with 1:500 goat anti-UIS4 (Sicgen, Cantanhede, Portugal) overnight at 4°C and 1:100 Green donkey anti-goat secondary antibody (Abbkine, Wuhan, China) for 1 h at room temperature. After washing with PBS three times, the cells were labeled with 1:500 rabbit anti-CSP overnight at 4°C and 1:100 Red donkey anti-rabbit secondary antibody (Abbkine) for 1 h at room temperature. The nuclei were counterstained with DAPI (Beyotime) for 5 min at room temperature.

For invasion rate observation, images were obtained at 6 h after infection, UIS4 staining were used to recognize the parasites in HepG2 cells. For evaluating the co-localization of CSP_mut_ and LC3, the cells were transfected with hLC3B-RFP plasmid and infected with CSP_wt_ or CSP_mut_ sporozoites, and then labeled with goat anti-UIS4 and the secondary antibody as stated above. To observe the effect of CSP on the IFN-γ-mediated killing of EEFs, control and CSP-stable transfected HepG2 cells were pre-treated with 1 U/mL recombinant human IFN-γ (Peprotech, Rocky Hill, NJ, USA) or equivoluminal medium for 6 h. The cells were then infected with *P.b* ANKA or *P.b* ANKA-RFP as stated above and co-treated with IFN-γ 46 h hours after infection.

For evaluating the effect of autophagy regulation on IFN-γ-mediated killing, HepG2 cells were pre-treated with 0.2 U/mL, 0.5 U/mL, and 1 U/mL IFN-γ for 6 h, and then infected with *P.b* ANKA-RFP sporozoites at a ratio of 1:3 and co-treated with IFN-γ and 0.25 μg/mL rapamycin, 10 μM LY294002 (Cell Signaling Technology, Danvers, MA, USA), 100nM wortmannin (Sigma-Aldrich), 1μM TAK-243 (MCE, NJ, USA) or equivoluminal medium respectively for 24 and 46 h.

For examining effect of CSP overexpression on the co-localization of EEF and LC3, control and CSP-stable transfected HepG2 cells were transfected with hLC3B-RFP plasmid. Twenty-four hours after transfection, the cells were infected with *P.b* ANKA sporozoites as stated above for 24 h, and then treated as described above.

For autophagy flux observation, the cells were transfected with the pcDNA3.1 vector or pcDNA3.1-CSP plasmids as stated above for 24 h. The supernatant was then replaced with fresh culture medium and the cells were infected with adenovirus containing tandem GFP-RFP-LC3 structures (Hanbio, Shanghai, China). Since the cell culture was changed, the cells were treated with 0.25 μg/mL rapamycin (Cell Signaling Technology) or equivoluminal medium respectively for 24 h. The cells were immobilized, penetrated, and stained with DAPI. After washing with PBS, the slides of cells were mounted on a microscope slide using DAKO Fluorescence Mounting Medium (Agilent). LSM 780 NLO microscope systems and ZEN Imaging Software (Zeiss) were used for image acquisition and export. Image J software (NIH) was used for analyzing the number and size of EEFs.

### Total RNA extraction and real-time PCR

For parasite burden detection, the livers of mice were dissected at 46 h after infection of CSP_wt_ or CSP_mut_ sporozoites, and homogenized in 1.5 mL Trizol (Invitrogen), and HepG2 cells were collected 46 h after infection of CSP_wt_ or CSP_mut_ sporozoites and lysed by 1 mL Trizol (Invitrogen). 100μL liver homogenate or 1 mL of cell lysis were used for total RNA extraction using Trizol in accordance with the manufacturer’s instructions. cDNA was synthesized from equivalent total RNA using PrimeScript™ RT reagent Kit with gDNA Eraser (TAKARA) in accordance with the manufacturer’s instructions. The parasite load in the livers and HepG2 cells was evaluated by Taqman-PCR with primers and probes for 18S rRNA and *GAPDH* following the manufacturer’s instructions of Premix Ex Taq™ (Probe qPCR) (TAKARA). For the SYBR quantitative PCR assay, HepG2 cells in 6-well plates were lyzed by 1 mL Trizol. Total RNA was extracted and cDNA was synthesized as described above. The mRNA levels of *IFN-γ*, *iNOS, LC3B, ATG3, ATG5, ATG7, STUB1, SYVN1, CBL, SMURF1, PAFAH1B1,* and *NEDD4* were evaluated using TB Green™ Premix Ex Taq™ II (Tli RNaseH Plus) (TAKARA). CFX96 Touch™ Real-Time PCR Detection System and CFX ManagerTM Software (Bio-Rad, Hercules, CA, USA) were used for RT-PCR data collection and analysis.

### Western blot and immunoprecipitation

To detect the levels of autophagy-related genes, 1 × 10^6^ control and CSP-stable transfected HepG2 cells were incubated in 6-well plates and treated with 0.25 μg/mL rapamycin for 0 h, 6 h, 12 h and 24 h. For half-life detection, the CSP-stable transfected HepG2 and control cells were treated with 20 μM cycloheximide (CHX, Cell Signaling Technology) or 10 μM MG132 (Cell Signaling Technology) for 0 h, 6 h, 12 h and 24 h. For ubiquitination inhibiting experiment, CSP-stable transfected HepG2 and control cells were treated with IFN-γ, 1μM TAK-243 or both for 24 h. The cells were lyzed with 300 μL RIPA lysis buffer containing a protease and phosphatase inhibitor cocktail (Thermo Fisher) at various time points and protein concentrations were detected by BCA Protein Assay Kit (Sangon Biotech). Total proteins (20 μg equivalent) were then separated by 10% or 12% sodium dodecyl sulfate-polyacrylamide gel electrophoresis (SDS-PAGE; Bio-Rad), then transferred by using 0.22-μm polyvinylidene fluoride (PVDF) filter (Milipore) and blocked with blocking solution (Sangon Biotech). The PVDF filter was incubated with 1:2000 rabbit anti-LC3B (Sigma-Aldrich), 1:2000 rabbit anti-ATG3 (Abcam, Cambridge, UK), 1:2000 rabbit anti-ATG5 (Invitrogen), 1:2000 rabbit anti-ATG7 (Sigma-Aldrich), 1:2000 rabbit anti-P62 (Sigma-Aldrich), 1:2000 rabbit anti-Beclin-1 (Cell Signaling Technology), and 1:2000 mouse anti-β-actin (Sigma-Aldrich) overnight at 4°C. Then, the PVDF filter was washed three times with TBS solution containing 1% Tween-20 (Amersco) and incubated with 1:20,000 goat anti-rabbit or anti-mouse IgG-HRP secondary antibody (Santa Cruz) for 1 h at room temperature.

For co-immunoprecipitation, 1 × 10^6^ CSP-stable transfected HepG2 and control cells in 6-well plates were transfected with 3 μg pFLAG-cmv8 or pFLAG-cmv8-human ATG5/7 and 1μg His-UB plasmids, respectively, and treated with 10 μM MG-132 for 24 h. 2 × 10^6^ 293FT cells in two 6-well plates were co-transfected with 4 μg pFLAG-cmv8-human ATG5/7 or 4 μg pcDNA3.1, pcDNA3.1-CSP, pcDNA3.1-CSPΔNLS, or pcDNA3.1-NLS and 2μg His-UB plasmids and treated with 10 μM MG-132 for 24 h. Then cells were lysed with 1 mL Western and IP Cell lysis Buffer. The protein concentrations of each group were detected and equalized as stated above. Fusion FLAG-ATG5 and ATG7 proteins were captured using Anti-FLAG^®^ M2 Magnetic Beads (Sigma-Aldrich) according to the manufacturer's instructions. After washing five times with equilibrium buffer (50 mM Tris HCl, 150 mM NaCl, pH 7.4), the beads were boiled with SDS-PAGE loading buffer for 10 min. Protein solution (20 μL equivoluminal) was loaded and separated by 10% SDS-PAGE as stated above. The PVDF filter was incubated with 1:500 mouse anti-DYKDDDDK Tag (Invitrogen), 1:1000 rabbit monoclonal anti-His-Tag (Cell Signaling Technology), 1:2000 mouse anti-β-actin (Sigma-Aldrich) or 1:500 rabbit anti-NEDD4 overnight at 4°C, and incubated with secondary antibody (Santa Cruz) as stated above for 1 h at room temperature. The protein bands were visualized using the Western BLoT Hyper HRP Substrate (TAKARA) and exposed using a Chemiluminescence Imaging System (Fusion Solo S, Vilber, France). Image J software (NIH) was used for grey value analysis.

### RNA interference

RNA interference for ATG5 and ATG7 in HepG2 cells was performed according to the protocol of ATG5/7 Human shRNA Plasmid Kits (Origene, Rockville, MD, USA). HepG2 cells (1 × 10^6^) in 6-well plates were transfected with 4 μg ATGs interfering or scrambled non-effective plasmids using Lipofectamine 3000 following the manufacturer protocol. The plasmids containing the following sequences were used for *ATG5* mRNA interference, 5′-TCAGCTCTTCCTTGGAACATCACAGTACA-3′, *ATG7* mRNA interference, 5′-CTTGGCTGCTACTTCTGCAATGATGTGGT-3′, and shNC sequence, 5′-GCACTACCAGAGCTAACTCAGATAGTACT-3′. For NEDD4 mRNA interference, 5′-GTGAAATTGCACATAATGAGGTTCAAGAGACCTCATTATGTGCAATTTCAC-3′, and shNC sequence, 5′-GTTCTCCGAACGTGTCACGTCAAGAGATTACGTGACACGTTC GGAGAA-3′. Fresh culture medium containing 3 mg/mL of puromycin (Gibco) was added to select puromycin-resistant cells. The RNA interference efficiency was validated by quantitative real-time PCR and western blot.

### Transcriptome sequencing

1 × 10^6^ CSP-stable transfected or control cells were lyzed by 1 mL Trizol (Invitrogen). Total RNA was extracted as stated above. A total amount of 2 μg RNA per sample was used as input material for the RNA sample preparations. mRNA was purified from total RNA using poly-T oligo-attached magnetic beads. Fragmentation was carried out using divalent cations under elevated temperature in VAHTSTM First Strand Synthesis Reaction Buffer (5X). First-strand cDNA was synthesized using a random hexamer primer and M-MuLV Reverse Transcriptase (RNase H). Second-strand cDNA synthesis was subsequently performed using DNA polymerase I and RNase H. Remaining overhangs were converted into blunt ends via exonuclease/polymerase activities. After adenylation of the 3′ ends of DNA fragments, an adaptor was ligated for library preparation. To select cDNA fragments of preferentially 150–200 bp in length, the library fragments were purified with the AMPure XP system (Beckman Coulter, Beverly, USA). Then, 3 μL USER Enzyme (NEB, Ipswich, MA, UK) was incubated with size-selected, adaptor-ligated cDNA at 37°C for 15 min followed by 5 min at 95°C before PCR. PCR was performed with Phusion High-Fidelity DNA polymerase, Universal PCR primers, and Index (X) Primer. Finally, PCR products were purified (AMPure XP system) and the library quality was assessed on the Agilent Bioanalyzer 2100 system. The libraries were then quantified and pooled. Paired-end sequencing of the library was performed on HiSeq XTen sequencers (Illumina, San Diego, CA). RAW data have been submitted to GEO database, and series record is GSE129323. Heatmap was generated by using the *R* language version 4.0.2 and the pheatmap package v1.0.12.

### Dual luciferase reporter assay

Promoter activity assay was conducted by co-transfecting 100ng pcDNA3.1 vector, pcDNA3.1-CSP, pcDNA3.1-CSPΔNLS or pcDNA3.1-NLS and 100ng pGL3-NEDD4 promoter and 1ng RL-TK into HEK293T cells in a 96-well, 24 h later, cells were lyzed and activities of luciferase for each group were detected by Dual-Luciferase® Reporter Assay System (Promega, Madison, WI, USA).

### Statistical analysis

SPSS 19.0 software (IBM, Armonk, NY, USA) was used for the statistical analysis. Student’s *t*-test for two groups or analysis of variance for multiple groups were used if the data were normally distributed for compare continuous variables, and if not, the Mann-Whitney U test was used for comparisons among groups. Pairwise differences in normally distributed variables were compared by the Tukey-Kramer statistic for multiple comparisons. A *p*-value of < 0.05 was considered statistically significant. Error bars represent standard errors of the mean.

## Acknowledgments

This work was supported by the National Natural Science Foundation of China (No. 81672053 and 81702247), the State Key Program of the National Natural Science Foundation of China (No. 81830067) and the Miaopu Talent Grant from Army Medical University (2019R057).

## Author contributions

Conceptualization, W.X., Y.W., H.Z., and J.D.; Writing – Original Draft, W.X. and H. Z; Writing–Review & Editing, W.X.; Y.W., J.D., and H. Z.; Methodology, H. Z., and L.X.; Investigation, H. Z., X.L., K.L., C.Z., T.L., F. Z., T.Z. Y. D., and Y. F.; Funding Acquisition, X.W.; Resources, W.X.; Y.W., and J. D.; Supervision, W.X., Y.W., and J.D.

## Conflict of interest

The authors declare no competing interests.

## Supplemental material

### Supplementary Figures

**Figure S1.**
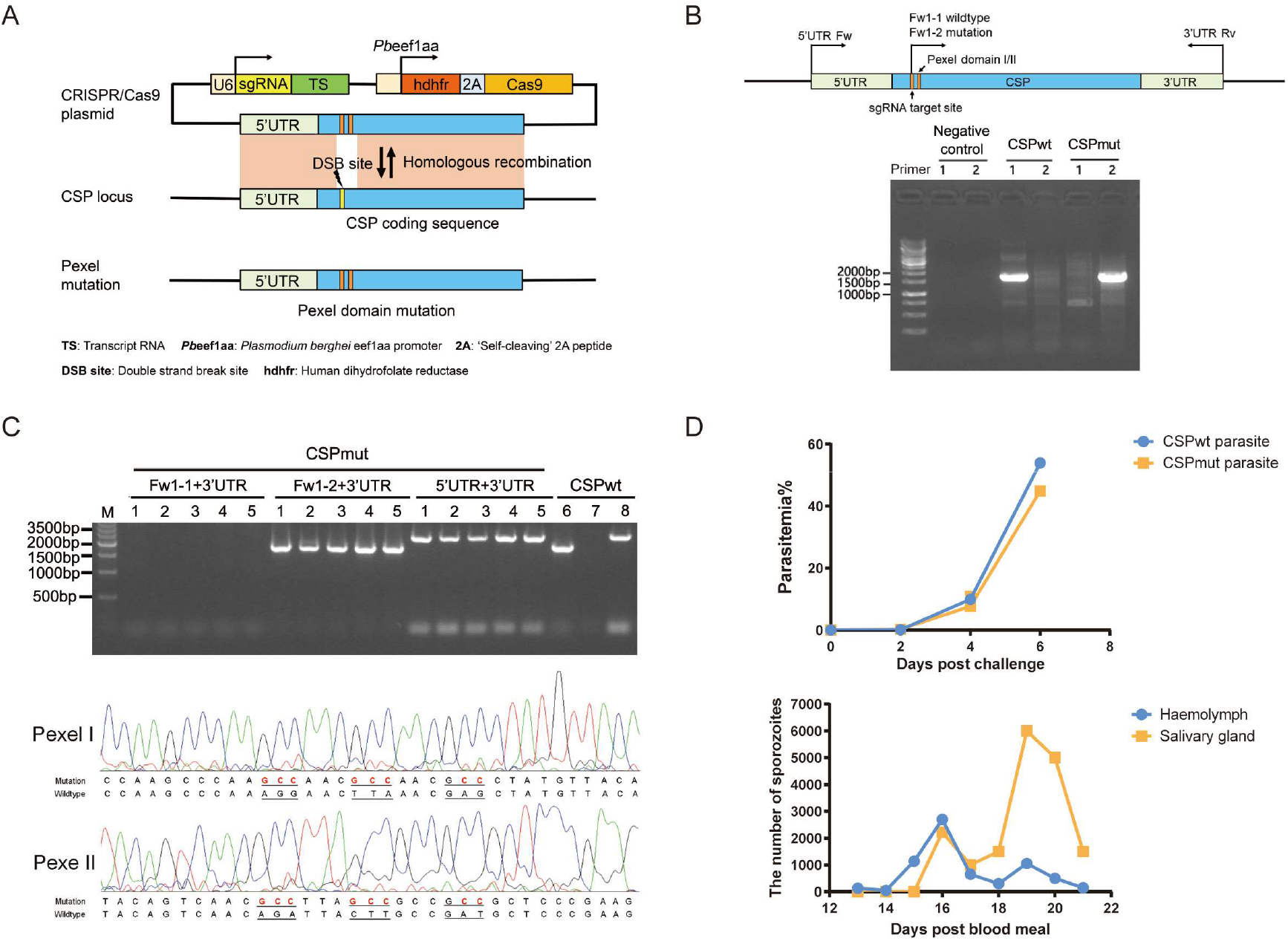
Construction and characterization of the CSP_mut_ parasite. (**A**) Schematic construction of the CSP_mut_ parasite by CRISPR-Cas9. The plasmid contains Cas9 and sgRNA expression cassettes and donor template pexel/II mutant CSP for homologous recombination repair after a double-strand break (DSB) at the WT CSP. (**B**) Primers designed to identify the WT and mutant parasite. A ~1.6-kb fragment could be amplified from the genomic DNA of the mutant parasite with Primer 2 (Fw1-2 (Mut) and 3′-UTR Rv), but no fragment was obtained with Primer 1 (Fw1-1 (WT) and 3′-UTR Rv). (**C**) Identification of CSP_mut_ parasite clones. The pyrimethamine-resistant parasites in mice were collected and cloned by injecting each mouse with ~1.0 infected iRBCs. The resulting clones were identified by PCR (*top*) and the *CSP* gene was sequenced (*bottom*). (**D**) The parasitemia of mice infected with the CSP_mut_ or CSP_wt_ parasite was determined (*top*), sporozoites were examined in both the hemolymph and salivary gland of mosquitoes at indicated times post-infection with CSP_mut_ parasite (*bottom*).

**Figure S2.**
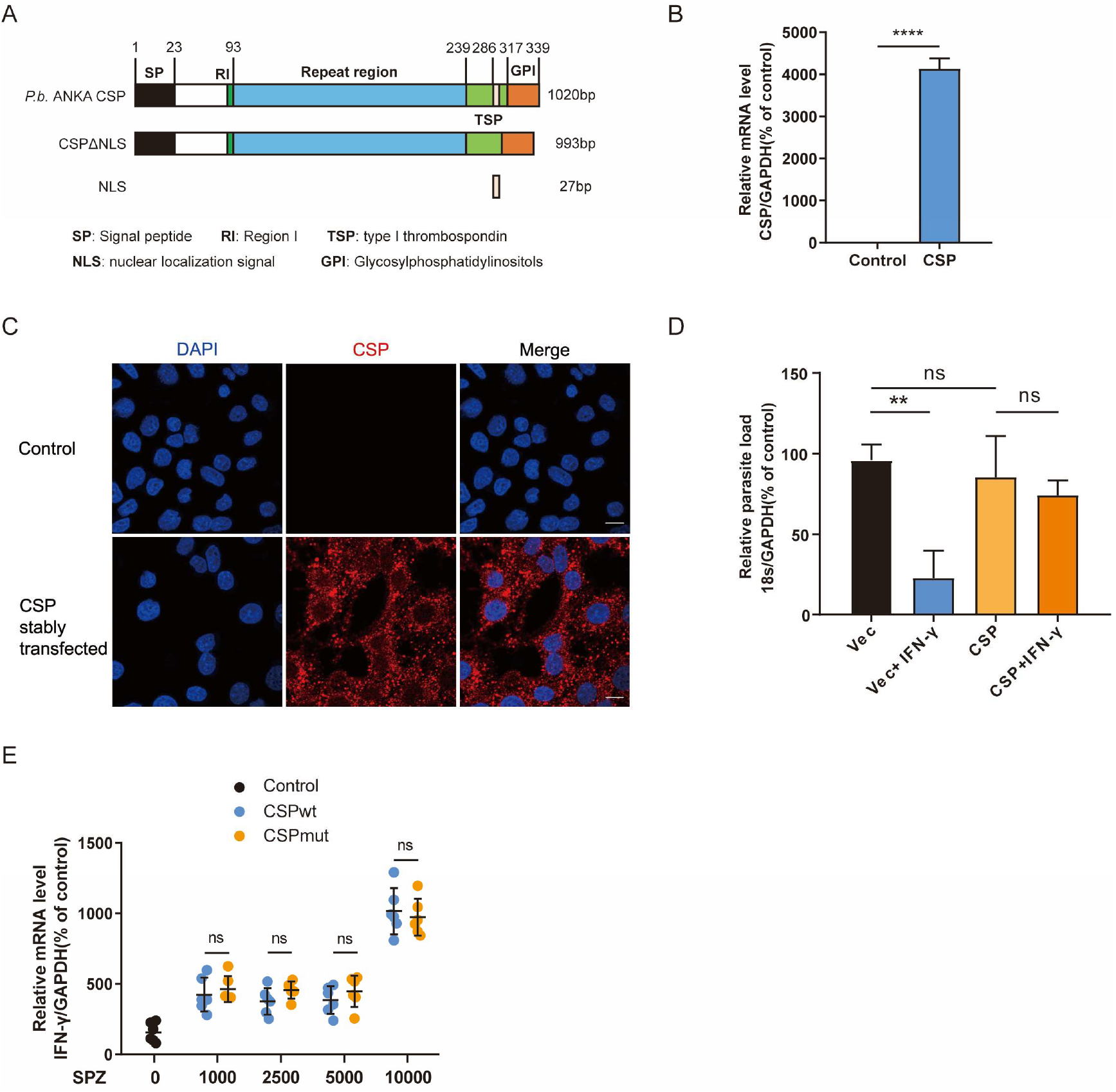
Schematic view of CSP protein and construction of CSP-stable transfected HepG2 cell line. (**A**) Schematic view of *P.b* ANKA CSP (Uniprot Entry: P06915), CSPΔNLS and NLS used in this study. **SP**, signal peptide; **RI**: Region I; **TSP**: type I thrombospondin; **NLS**, nuclear localization signal. **GPI**, glycosylphosphatidyl inositol attachment site. (**B**) CSP expression detected by real-time qPCR in control and CSP-stable transfected HepG2 cells. (**C**) Indirect immunofluorescence assay showed the distribution of CSP protein in cytoplasm CSP-stable transfected HepG2 cell, CSP protein was labeled by Poly rabbit anti-CSP antibody (1:200) and IFKine Red AffiniPure donkey anti-rabbit IgG (H+L) (1:500). Scale bar=10 μm. (**D**) Control and CSP-stable transfected HepG2 cells were pre-treated with or without IFN-γ followed by incubation with CSP_wt_ or CSP_mut_ sporozoites; 46 h later, the parasite burden was determined as described as above. (**E**) mRNA levels of IFN-γ were evaluated in the mouse liver 46 h after infection of indicated amount of CSP_wt_ or CSP_mut_ sporozoites by real-time PCR (Control indicated non-infected mice). Data are represented as mean ± SEM; ns, not significant; \*\*\*\**p*<0.0001.

**Figure S3.**
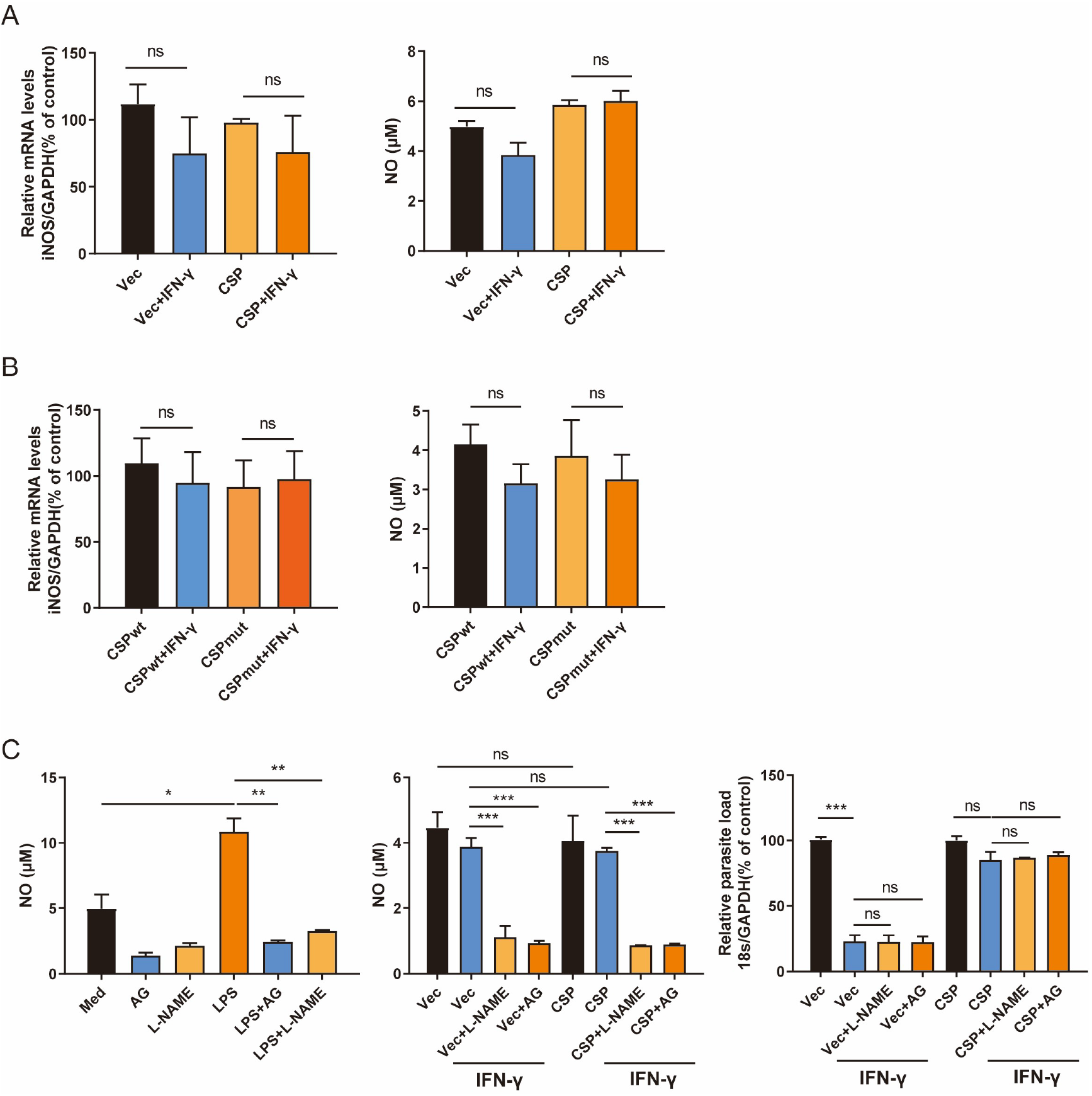
NO is not involved in the suppression of the IFN-γ-mediated suppression of EEFs by CSP. (**A**) 1.2 × 10^5^ HepG2 cells were transfected with pcDNA3.1 (Vec) or pcDNA3.1-CSP plasmid and treated with or without IFN-γ, and then infected with 4 × 10^4^ *P.b* ANKA sporozoites for 24 h. The mRNA level of iNOS (*left*) and the concentration of NO in the supernatants (*right*) were measured by real-time qPCR and the Griess reaction assay, respectively. (**B**) 1.2 × 10^5^ HepG2 cells were pre-treated with or without IFN-γ and incubated with 4 × 10^4^ WT or CSP_mut_ sporozoites for 24 h. The mRNA level of iNOS (*left*) and concentration of NO (*right*) produced by CSP_wt_ or CSP_mut_ parasite-infected hepatocytes were measured as described as above. (**C**) RAW264.7 cells were pretreated with or without the iNOS inhibitor Aminoguanidine (AG) or L-NAME and then stimulated with LPS for 4 h. The concentration of NO in the supernatants of LPS-treated macrophages was measured (*left*). 1.2 × 10^5^ HepG2 cells were transfected with pcDNA3.1 (Vec) or pcDNA3.1-CSP plasmid and treated with or without the iNOS inhibitor AG or L-NAME and IFN-γ followed by infection with 4 × 10^4^ *P.b* ANKA sporozoites. The concentration of NO (*middle*) and the parasite load (*right*) were determined by 24 h and 46 h after infection, respectively. Data are represented as mean ± SEM; ns, not significant; \**p* < 0.05, \*\**p* < 0.01, \*\*\**p* < 0.001.

**Figure S4.**
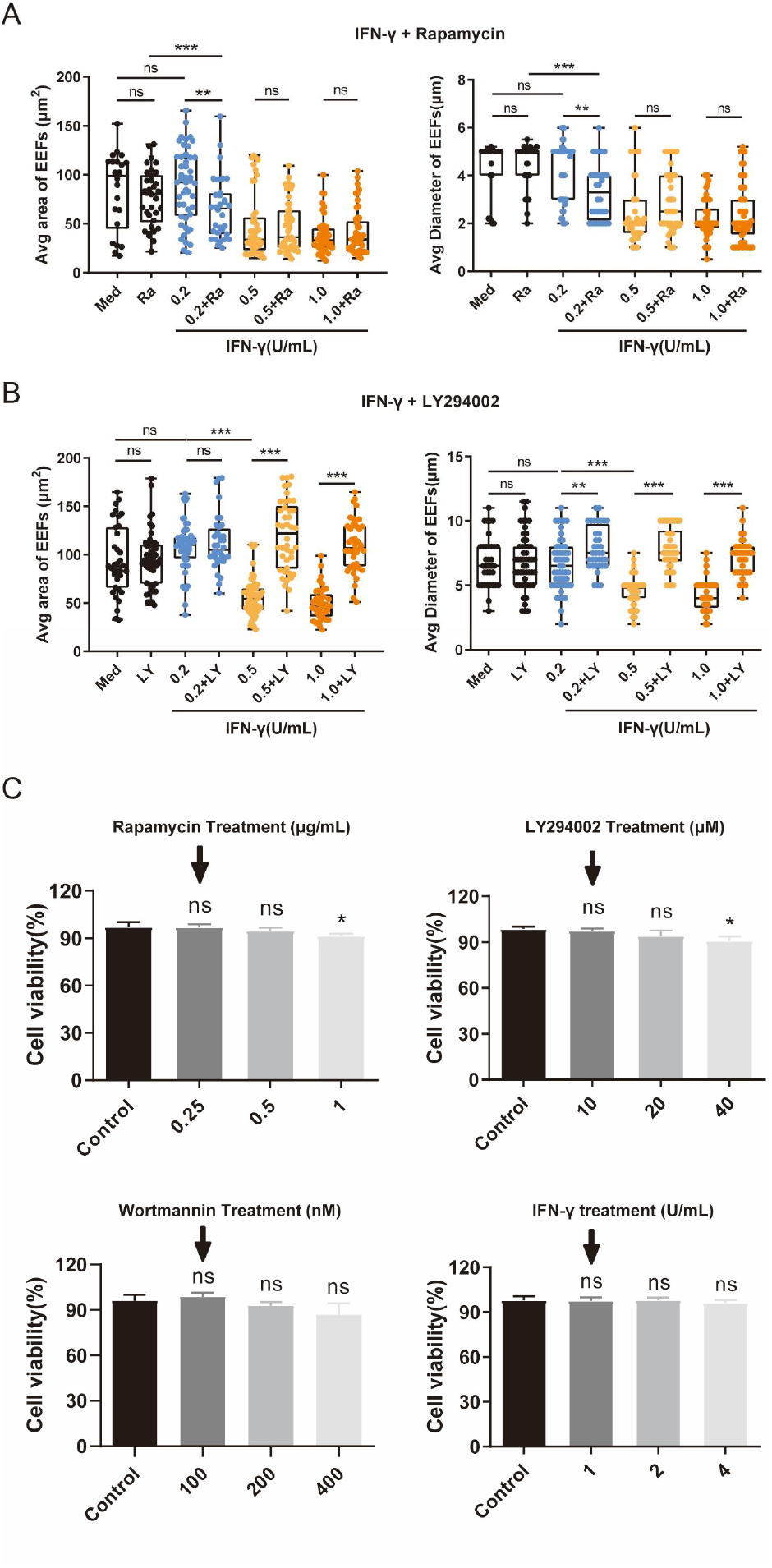
The effect of the autophagy modulators on the IFN-γ-suppression of EFFs development. (**A**) Effect of the induction of autophagy on the IFN-γ-mediated suppression of EEFs *in vitro*. 1.2 × 10^5^ HepG2 cells were pre-treated with or without the autophagy inducer rapamycin (Rapa) and IFN-γ at the indicated concentrations and then incubated with 4 × 10^4^ sporozoites. The size (*left*) and the diameter (*right*) of EEFs at 24 h post-infection was compared. (**B**) Effect of the inhibition of autophagy on IFN-γ-mediated suppression of EEFs *in vitro*. HepG2 cells were pre-treated with or without the autophagy inhibitor LY294002 (LY) and IFN-γ at the indicated concentrations and then incubated with sporozoites. The size (*left*) and diameter (*right*) of EEFs at 24 h post-infection was compared. (**C**) CCK-8 analysis results showed that chemical agents used as mentioned above in these experiments have no influences on the cell viability of HepG2 cells (Black arrow indicates the concentration of chemical agents used in our study). Data are represented as mean ± SEM; ns, not significant; *p < 0.05; **p<0.01; ***p<0.001.

**Figure S5.**
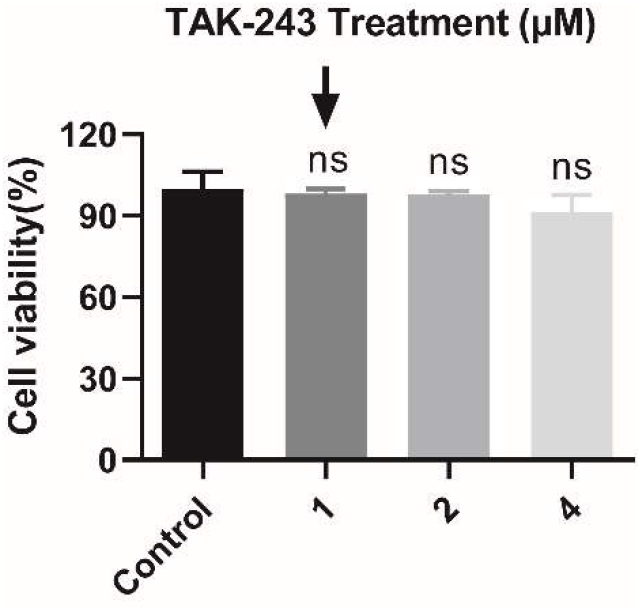
CCK-8 analysis showed that TAK-243 used in our experiments have no influences on the cell viability of HepG2 cells. (Black arrow indicates the concentration of TAK-243 used in our study). Data are represented as mean ± SEM; ns, not significant.

**Figure S6.**
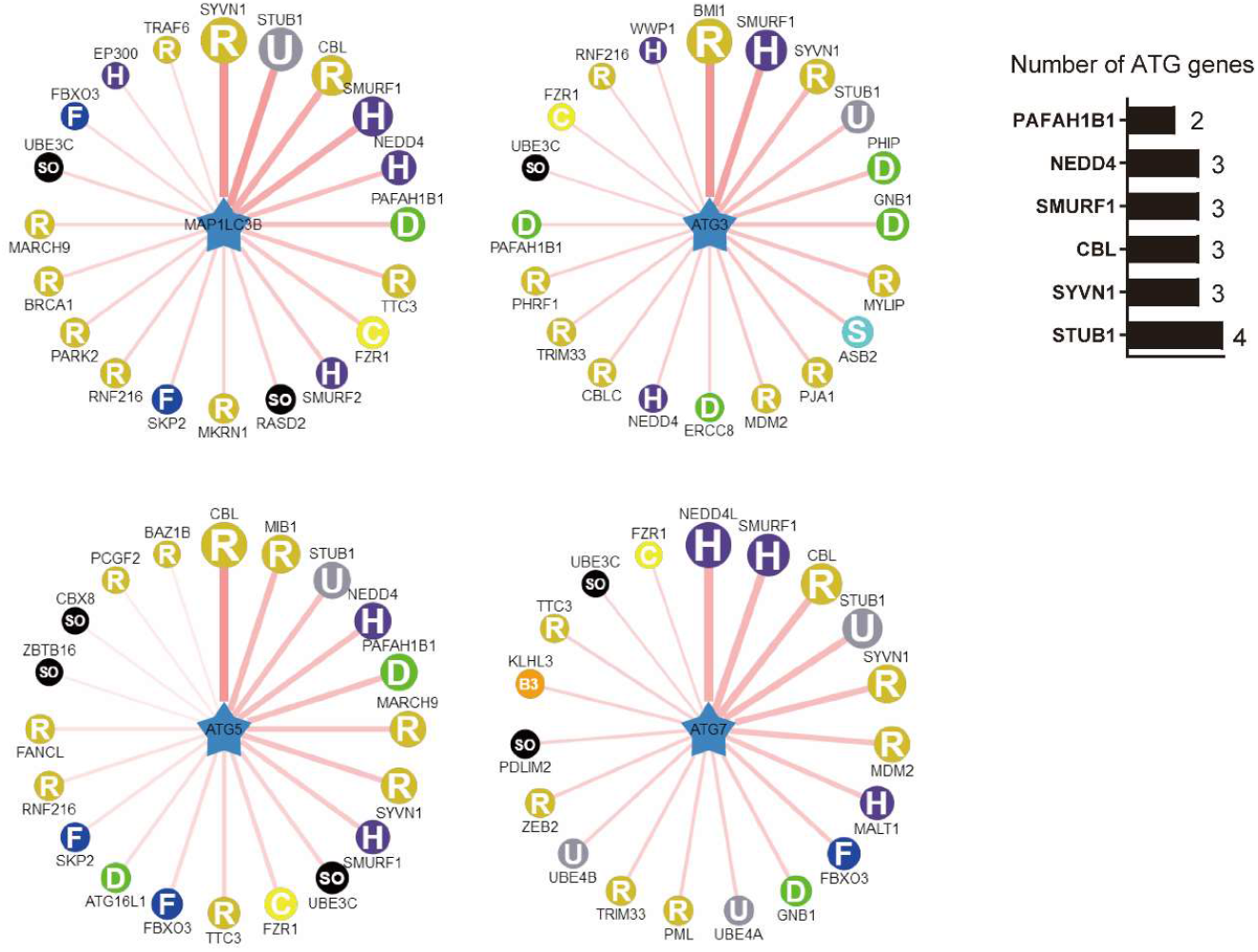
The prediction of E3 ubiquitin ligases possibly involved in the ubiquitination of ATGs. E3 ubiquitin ligases possibly involved in the ubiquitination of ATGs were predicted on line (http://ubibrowser.ncpsb.org/ubibrowser/) (*left*), and E3 ubiquitin ligases predicted to regulate at least two ATGs (LC3, ATG3, ATG5 and ATG7) were listed in the right column. Numbers indicated the number of ATG was regulated by the six predicted E3 ubiquitin ligases, including STUB1, SYVN1, CBL, SMURF1, PAFAH1B1 and NEDD4, respectively. (*right*).

**Figure S7.**
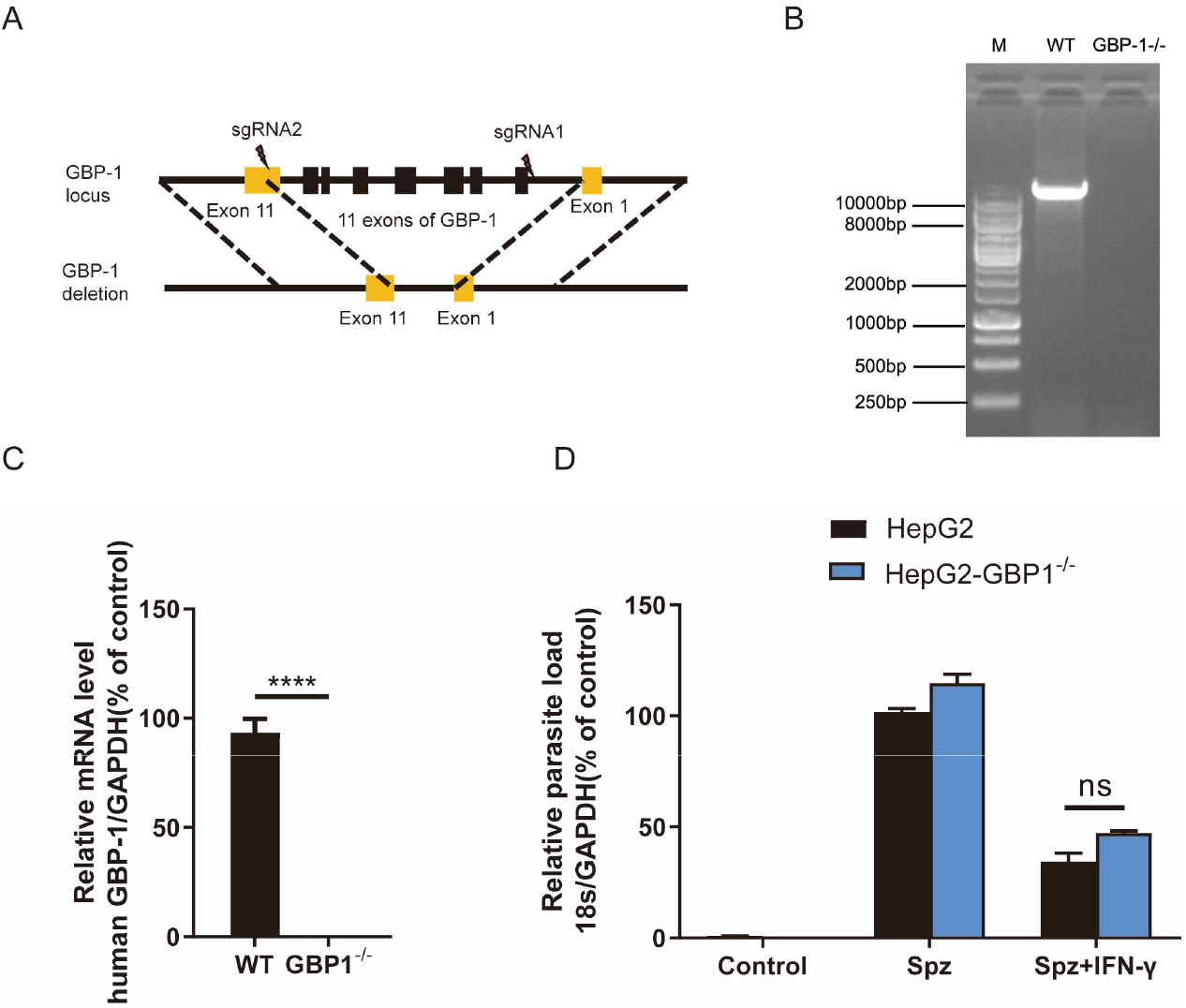
GBP1 is dispensable for the IFN-γ-mediated killing of EEFs. (**A**) Schematic representation of the disruption of *GBP1* gene by CRISPR-Cas9. (**B**) The identification of *GBP1*^−/−^ HepG2 cells by PCR with specific primers binding to the left side of the sgRNA2 and right side of sgRNA1 targeting site in *GBP-1* exons (PCR fragment of wide type is 11,448bp). (**C**) The relative mRNA level of *GBP1* to *GAPDH* was detected in WT and *GBP1*^−/−^ HepG2 cells by real-time PCR. (**D**) 1.2 × 10^5^ WT and *GBP1*^−/−^ HepG2 cells were infected with 4 × 10^4^ *P.b* ANKA sporozoites for 46 h, and then treated with IFN-γ. The parasite load in WT and *GBP1*^−/−^ HepG2 cells was measured and compared as described before. Data are represented as mean ± SEM; \*\*\*\**p* < 0.0001; ns, not significant.

### Supplementary Tables

**Table S1.**
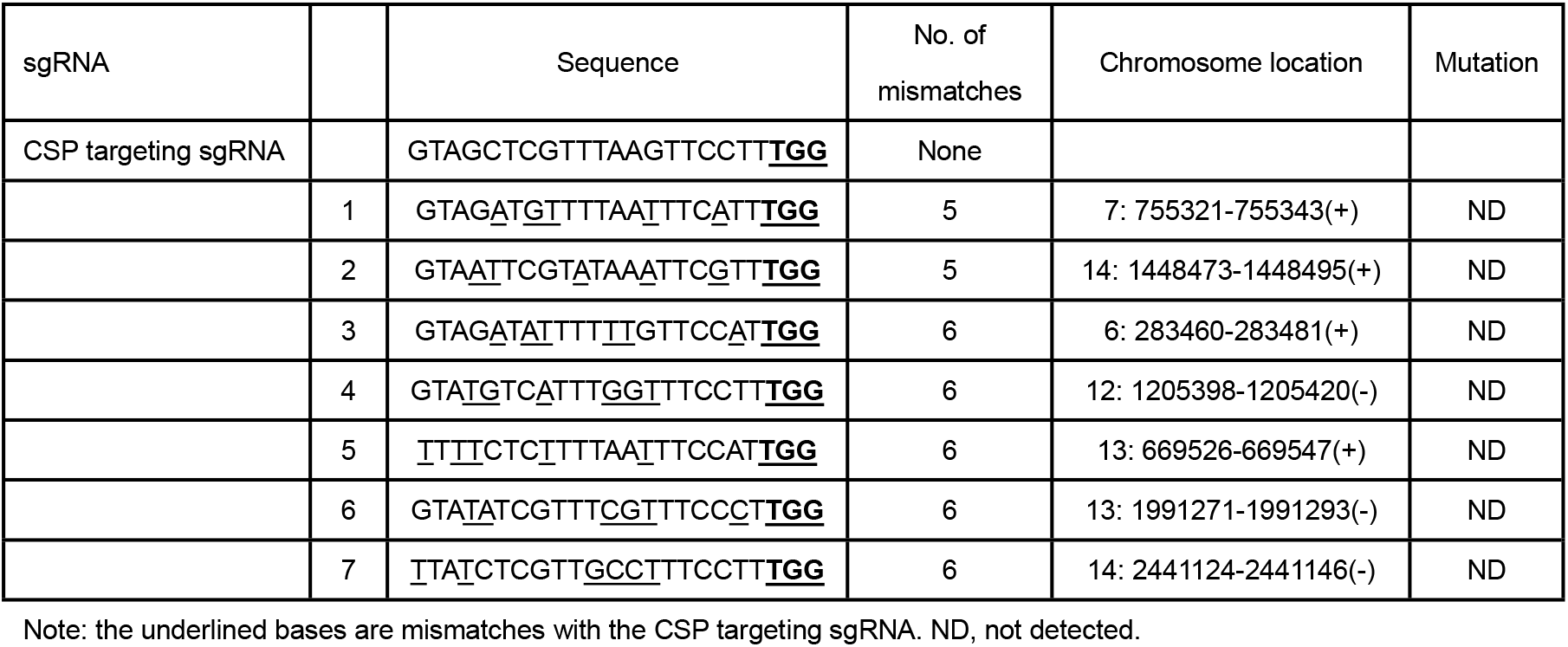
Numbers of mutations detected at the potential off-target cleavage sites in the *Plasmodium berghei* ANKA genome.

**Table S2.** Scoring of E3 ubiquitin ligases possibly involved in the ubiquitination of each ATGs.

| **LC3** |  |  |  |  |  |  |  |  |  |
| --- | --- | --- | --- | --- | --- | --- | --- | --- | --- |
| E3 | E3GENE | SUB | SUBGENE | HOMO | PFAM | GO | NET | MOTIF | SCORE |
| Q86TM6 | SYVN1 | Q9GZQ8 | MAP1LC3B | 1 | 1 | 1.25 | 1 | 6.61 | 0.714 |
| Q9UNE7 | STUB1 | Q9GZQ8 | MAP1LC3B | 1 | 1 | 3.77 | 1.84 | 1 | 0.699 |
| P22681 | CBL | Q9GZQ8 | MAP1LC3B | 1 | 1 | 3.77 | 1.44 | 1 | 0.676 |
| Q9HCE7 | SMURF1 | Q9GZQ8 | MAP1LC3B | 1 | 1 | 2.88 | 2.3 | 1.06 | 0.675 |
| P43034 | PAFAH1B1 | Q9GZQ8 | MAP1LC3B | 1 | 1 | 3.77 | 1 | 1 | 0.64 |
| P46934 | NEDD4 | Q9GZQ8 | MAP1LC3B | 1 | 1 | 3.77 | 1 | 1 | 0.64 |
| P53804 | TTC3 | Q9GZQ8 | MAP1LC3B | 1 | 1 | 1.25 | 1.29 | 2.12 | 0.63 |
| Q9UM11 | FZR1 | Q9GZQ8 | MAP1LC3B | 1 | 1 | 1.51 | 1 | 2.12 | 0.624 |
| Q9HAU4 | SMURF2 | Q9GZQ8 | MAP1LC3B | 1 | 1 | 1.78 | 1.77 | 1 | 0.622 |
| Q96D21 | RASD2 | Q9GZQ8 | MAP1LC3B | 1 | 1 | 1.51 | 1.84 | 1 | 0.609 |
| Q9NWF9 | RNF216 | Q9GZQ8 | MAP1LC3B | 1 | 1 | 1 | 1.29 | 2.12 | 0.608 |
| Q9UHC7 | MKRN1 | Q9GZQ8 | MAP1LC3B | 1 | 1 | 1 | 1.29 | 2.12 | 0.608 |
| Q13309 | SKP2 | Q9GZQ8 | MAP1LC3B | 1 | 1 | 1 | 1.29 | 2.12 | 0.608 |
| P38398 | BRCA1 | Q9GZQ8 | MAP1LC3B | 1 | 1 | 1.51 | 1.77 | 1 | 0.605 |
| O60260 | PARK2 | Q9GZQ8 | MAP1LC3B | 1 | 1 | 1.51 | 2.3 | 1 | 0.605 |
| Q86YJ5 | 9-Mar | Q9GZQ8 | MAP1LC3B | 1 | 1 | 1.25 | 1 | 2.12 | 0.604 |
| Q15386 | UBE3C | Q9GZQ8 | MAP1LC3B | 1 | 1 | 1.25 | 1 | 2.12 | 0.604 |
| Q9UK99 | FBXO3 | Q9GZQ8 | MAP1LC3B | 1 | 1 | 1.13 | 1 | 2.12 | 0.594 |
| Q09472 | EP300 | Q9GZQ8 | MAP1LC3B | 1 | 1 | 1.25 | 1.84 | 1 | 0.589 |
| Q9Y4K3 | TRAF6 | Q9GZQ8 | MAP1LC3B | 1 | 1 | 1.25 | 2.39 | 1 | 0.589 |
| **ATG3** |  |  |  |  |  |  |  |  |  |
| E3 | E3GENE | SUB | SUBGENE | HOMO | PFAM | GO | NET | MOTIF | SCORE |
| P35226 | BMI1 | Q9NT62 | ATG3 | 1 | 1 | 1.25 | 1 | 6.61 | 0.714 |
| Q9HCE7 | SMURF1 | Q9NT62 | ATG3 | 1 | 1 | 4.05 | 1.44 | 1.06 | 0.688 |
| Q86TM6 | SYVN1 | Q9NT62 | ATG3 | 1 | 1 | 1.25 | 1 | 3.41 | 0.652 |
| Q9UNE7 | STUB1 | Q9NT62 | ATG3 | 1 | 1 | 3.77 | 1 | 1 | 0.64 |
| Q8WWQ0 | PHIP | Q9NT62 | ATG3 | 1 | 1 | 2.88 | 1.29 | 1 | 0.639 |
| P62873 | GNB1 | Q9NT62 | ATG3 | 1 | 1 | 2.88 | 1.29 | 1 | 0.639 |
| Q8WY64 | MYLIP | Q9NT62 | ATG3 | 1 | 1 | 1.25 | 1 | 2.8 | 0.633 |
| Q96Q27 | ASB2 | Q9NT62 | ATG3 | 1 | 1 | 1 | 1 | 3.41 | 0.63 |
| Q8NG27 | PJA1 | Q9NT62 | ATG3 | 1 | 1 | 1.13 | 1 | 2.8 | 0.623 |
| Q00987 | MDM2 | Q9NT62 | ATG3 | 1 | 1 | 2.88 | 1 | 1.06 | 0.619 |
| Q13216 | ERCC8 | Q9NT62 | ATG3 | 1 | 1 | 2.88 | 1 | 1 | 0.613 |
| Q9P1Y6 | PHRF1 | Q9NT62 | ATG3 | 1 | 1 | 2.88 | 1 | 1 | 0.613 |
| Q9ULV8 | CBLC | Q9NT62 | ATG3 | 1 | 1 | 2.88 | 1 | 1 | 0.613 |
| P43034 | PAFAH1B1 | Q9NT62 | ATG3 | 1 | 1 | 2.88 | 1 | 1 | 0.613 |
| P46934 | NEDD4 | Q9NT62 | ATG3 | 1 | 1 | 2.88 | 1 | 1 | 0.613 |
| Q9UPN9 | TRIM33 | Q9NT62 | ATG3 | 1 | 1 | 2.88 | 1 | 1 | 0.613 |

| Q9H0M0 | WWP1 | Q9NT62 | ATG3 | 1 | 1 | 1.25 | 1 | 2.12 | 0.604 |
| --- | --- | --- | --- | --- | --- | --- | --- | --- | --- |
| Q9NWF9 | RNF216 | Q9NT62 | ATG3 | 1 | 1 | 1.25 | 1 | 2.12 | 0.604 |
| Q5XUX0 | FBXO31 | Q9NT62 | ATG3 | 1 | 1 | 1.25 | 1 | 2.12 | 0.604 |
| Q15386 | UBE3C | Q9NT62 | ATG3 | 1 | 1 | 1.25 | 1 | 2.12 | 0.604 |
| Q9UM11 | FZR1 | Q9NT62 | ATG3 | 1 | 1 | 1.25 | 1 | 2.12 | 0.604 |
| Q14258 | TRIM25 | Q9NT62 | ATG3 | 1 | 1 | 1.25 | 1 | 2.12 | 0.604 |
| Q9C040 | TRIM2 | Q9NT62 | ATG3 | 1 | 1 | 1.25 | 1 | 2.12 | 0.604 |
| **ATG5** |  |  |  |  |  |  |  |  |  |
| E3 | E3GENE | SUB | SUBGENE | HOMO | PFAM | GO | NET | MOTIF | SCORE |
| P22681 | CBL | Q9H1Y0 | ATG5 | 1 | 1 | 3.77 | 1 | 2.12 | 0.711 |
| Q86YT6 | MIB1 | Q9H1Y0 | ATG5 | 1 | 1 | 1.51 | 1 | 2.8 | 0.652 |
| Q9UNE7 | STUB1 | Q9H1Y0 | ATG5 | 1 | 1 | 3.77 | 1 | 1 | 0.64 |
| P46934 | NEDD4 | Q9H1Y0 | ATG5 | 1 | 1 | 3.77 | 1 | 1 | 0.64 |
| P43034 | PAFAH1B1 | Q9H1Y0 | ATG5 | 1 | 1 | 3.77 | 1 | 1 | 0.64 |
| Q86YJ5 | MARCH9 | Q9H1Y0 | ATG5 | 1 | 1 | 1.25 | 1 | 2.8 | 0.633 |
| Q86TM6 | SYVN1 | Q9H1Y0 | ATG5 | 1 | 1 | 1.25 | 1 | 2.8 | 0.633 |
| Q9HCE7 | SMURF1 | Q9H1Y0 | ATG5 | 1 | 1 | 2.88 | 1 | 1.06 | 0.619 |
| Q9UM11 | FZR1 | Q9H1Y0 | ATG5 | 1 | 1 | 1.25 | 1 | 2.12 | 0.604 |
| P53804 | TTC3 | Q9H1Y0 | ATG5 | 1 | 1 | 1.25 | 1 | 2.12 | 0.604 |
| Q15386 | UBE3C | Q9H1Y0 | ATG5 | 1 | 1 | 1.25 | 1 | 2.12 | 0.604 |
| Q9UK99 | FBXO3 | Q9H1Y0 | ATG5 | 1 | 1 | 1.13 | 1 | 2.12 | 0.594 |
| Q676U5 | ATG16L1 | Q9H1Y0 | ATG5 | 1 | 1 | 1.25 | 2.09 | 1 | 0.585 |
| Q9NWF9 | RNF216 | Q9H1Y0 | ATG5 | 1 | 1 | 1 | 1 | 2.12 | 0.581 |
| Q9NW38 | FANCL | Q9H1Y0 | ATG5 | 1 | 1 | 1 | 1 | 2.12 | 0.581 |
| Q13309 | SKP2 | Q9H1Y0 | ATG5 | 1 | 1 | 1 | 1 | 2.12 | 0.581 |
| Q05516 | ZBTB16 | Q9H1Y0 | ATG5 | 1 | 1 | 1.25 | 1.44 | 1 | 0.563 |
| Q9UIG0 | BAZ1B | Q9H1Y0 | ATG5 | 1 | 1 | 1.78 | 1 | 1 | 0.562 |
| P35227 | PCGF2 | Q9H1Y0 | ATG5 | 1 | 1 | 1.78 | 1 | 1 | 0.562 |
| Q9HC52 | CBX8 | Q9H1Y0 | ATG5 | 1 | 1 | 1.78 | 1 | 1 | 0.562 |
| **ATG7** |  |  |  |  |  |  |  |  |  |
| E3 | E3GENE | SUB | SUBGENE | HOMO | PFAM | GO | NET | MOTIF | SCORE |
| Q96PU5 | NEDD4L | O95352 | ATG7 | 1 | 1 | 2.33 | 1 | 2.12 | 0.667 |
| Q9HCE7 | SMURF1 | O95352 | ATG7 | 1 | 1 | 2.88 | 1.44 | 1.06 | 0.655 |
| P22681 | CBL | O95352 | ATG7 | 1 | 1 | 1.51 | 1 | 2.8 | 0.652 |
| Q9UNE7 | STUB1 | O95352 | ATG7 | 1 | 1 | 4.05 | 1 | 1 | 0.647 |
| Q86TM6 | SYVN1 | O95352 | ATG7 | 1 | 1 | 1.13 | 1 | 3.41 | 0.642 |
| Q00987 | MDM2 | O95352 | ATG7 | 1 | 1 | 1.27 | 1 | 2.8 | 0.634 |
| Q9UDY8 | MALT1 | O95352 | ATG7 | 1 | 1 | 2.33 | 1.44 | 1 | 0.628 |
| Q9UK99 | FBXO3 | O95352 | ATG7 | 1 | 1 | 1.13 | 1 | 2.8 | 0.623 |
| P62873 | GNB1 | O95352 | ATG7 | 1 | 1 | 2.33 | 1.29 | 1 | 0.617 |
| O60315 | ZEB2 | O95352 | ATG7 | 1 | 1 | 2.88 | 1 | 1 | 0.613 |
| Q14139 | UBE4A | O95352 | ATG7 | 1 | 1 | 2.88 | 1 | 1 | 0.613 |
| P29590 | PML | O95352 | ATG7 | 1 | 1 | 2.88 | 1 | 1 | 0.613 |
| O95155 | UBE4B | O95352 | ATG7 | 1 | 1 | 2.88 | 1 | 1 | 0.613 |
| Q9UPN9 | TRIM33 | O95352 | ATG7 | 1 | 1 | 2.88 | 1 | 1 | 0.613 |
| Q96JY6 | PDLIM2 | O95352 | ATG7 | 1 | 1 | 1 | 1 | 2.8 | 0.61 |
| P53804 | TTC3 | O95352 | ATG7 | 1 | 1 | 1 | 1 | 2.8 | 0.61 |
| Q9UH77 | KLHL3 | O95352 | ATG7 | 1 | 1 | 1 | 1 | 2.8 | 0.61 |
| Q15386 | UBE3C | O95352 | ATG7 | 1 | 1 | 1.25 | 1 | 2.12 | 0.604 |
| Q9UM11 | FZR1 | O95352 | ATG7 | 1 | 1 | 1.25 | 1 | 2.12 | 0.604 |

**Table S3:**
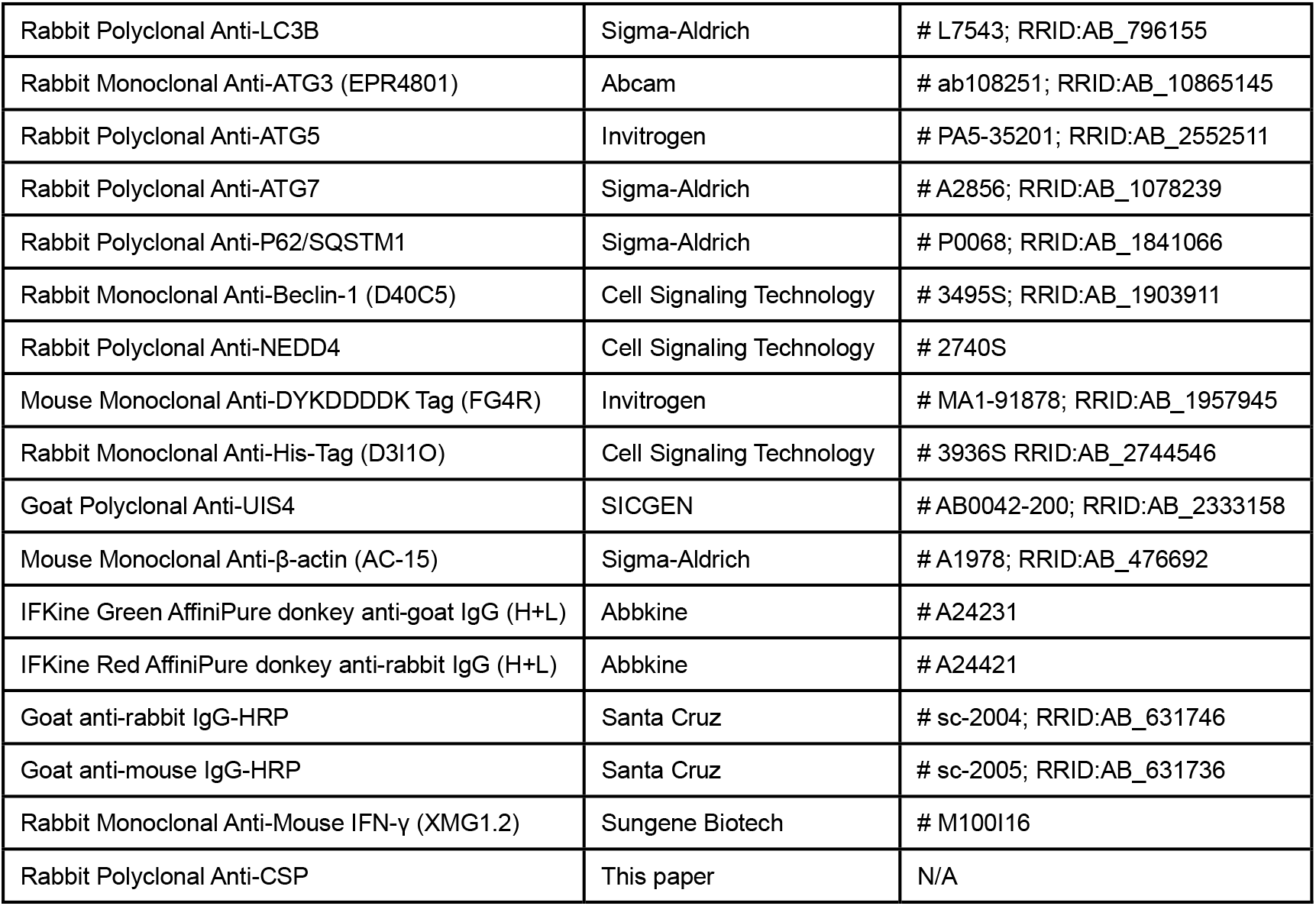
Antibodies used in this study.

**Table S4:**
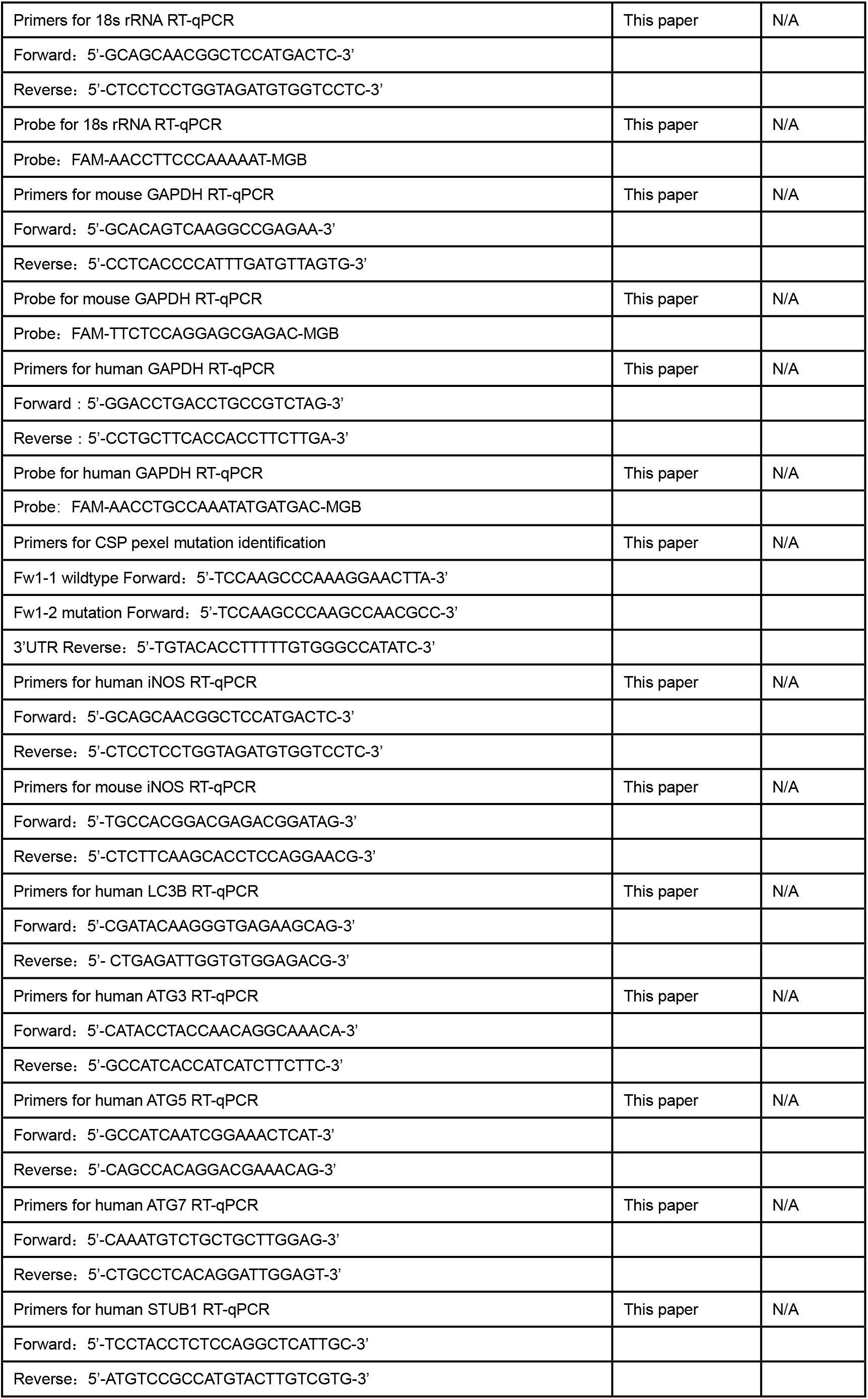
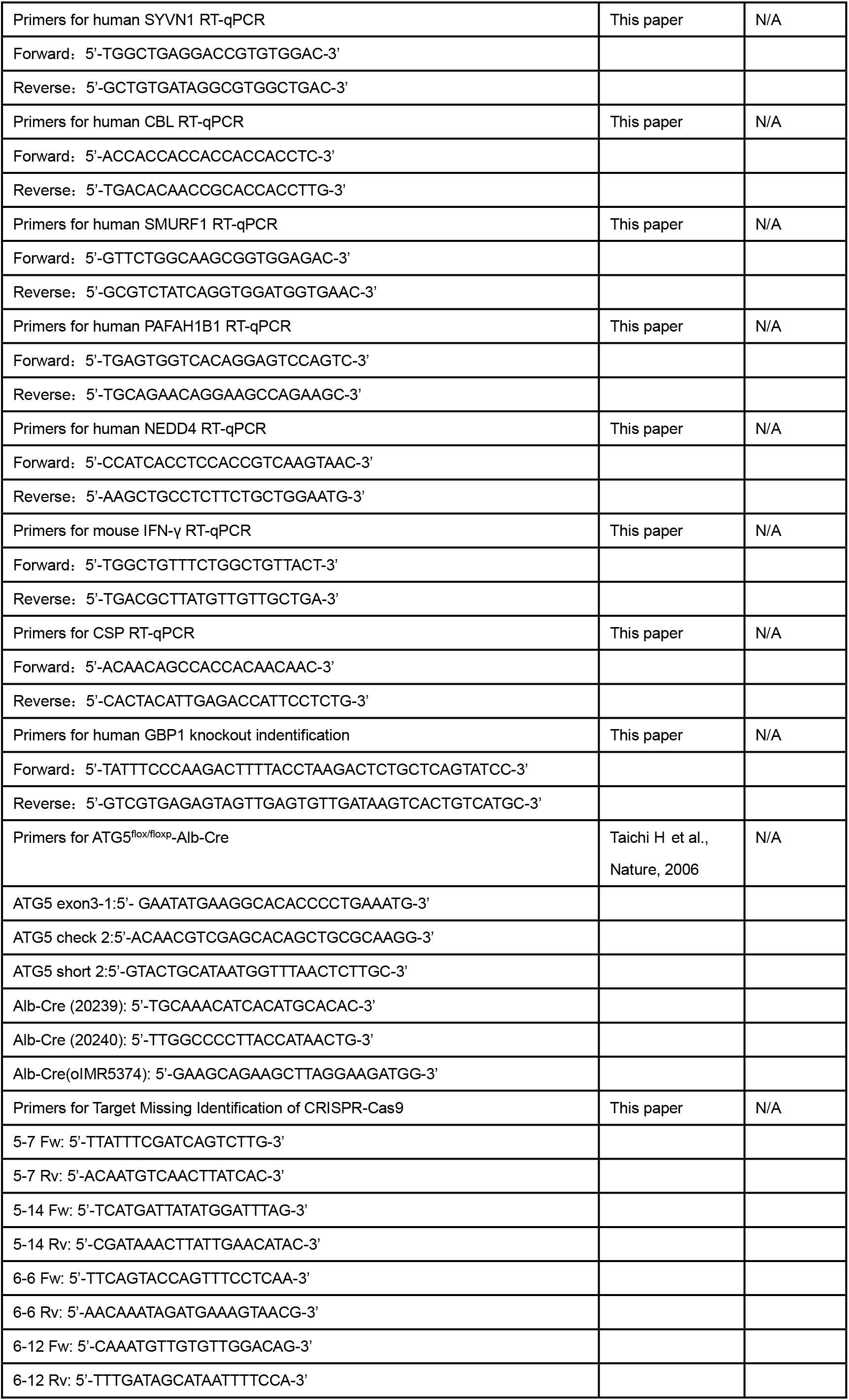
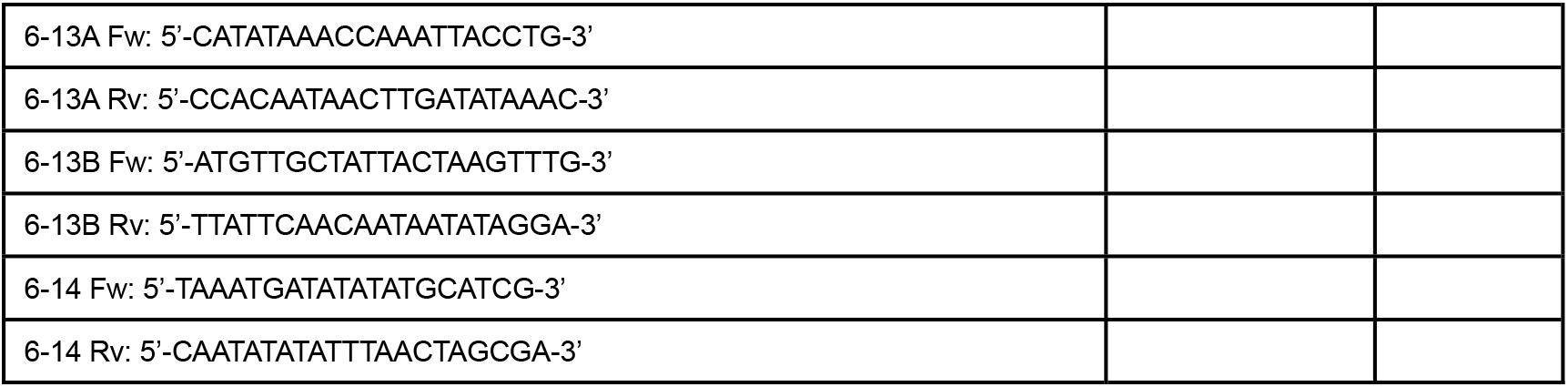
Oligonucleotides used in this study.

**Table S5:**
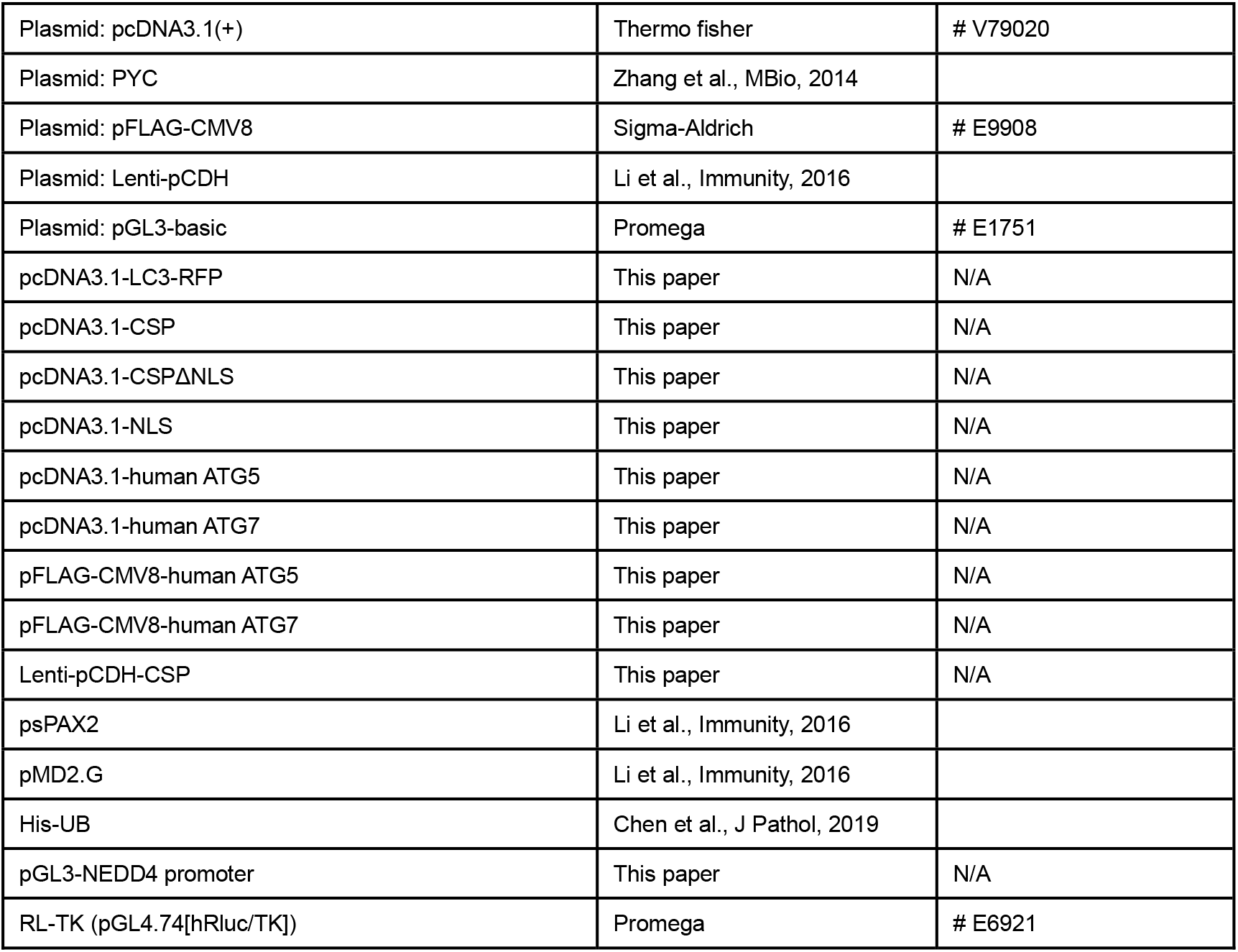
Plasmids used in this study.

|  |  |  |
| --- | --- | --- |
| Plasmid: pcDNA3.1(+) | Thermo fisher | # V79020 |
| Plasmid: PYC | Zhang et al., MBio, 2014 |  |
| Plasmid: pFLAG-CMV8 | Sigma-Aldrich | # E9908 |
| Plasmid: Lenti-pCDH | Li et al., Immunity, 2016 |  |
| Plasmid: pGL3-basic | Promega | # E1751 |
| pcDNA3.1-LC3-RFP | This paper | N/A |
| pcDNA3.1-CSP | This paper | N/A |
| pcDNA3.1-CSPΔNLS | This paper | N/A |
| pcDNA3.1-NLS | This paper | N/A |
| pcDNA3.1-human ATG5 | This paper | N/A |
| pcDNA3.1-human ATG7 | This paper | N/A |
| pFLAG-CMV8-human ATG5 | This paper | N/A |
| pFLAG-CMV8-human ATG7 | This paper | N/A |
| Lenti-pCDH-CSP | This paper | N/A |
| psPAX2 | Li et al., Immunity, 2016 |  |
| pMD2.G | Li et al., Immunity, 2016 |  |
| His-UB | Chen et al., J Pathol, 2019 |  |
| pGL3-NEDD4 promoter | This paper | N/A |
| RL-TK (pGL4.74[hRLuc/TK]) | Promega | # E6921 |

## Supplementary Materials and Methods

### Construction of CSP pexel I-II mutation parasite

The *P.b* ANKA CSP_mut_ parasite was constructed by replacement of the WT *CSP* sequence with a pexel I-II mutant *CSP* using CRISPR-Cas9. Single-guide RNA (5′-ATAGCTCGTTTAAGTTCCTT-3′) specifically targeting the *P.b.* ANKA *CSP* gene (Gene ID in PlasmoDB database PBANKA_0403200) was inserted downstream of the *Plasmodium* U6 promoter in the pYC plasmid (a gift from Dr. Jin Yuan, Xia’men University, China). The homologous recombinant fragment for the CSP pexel mutation containing the 5′ untranslated region (593 bp) and coding sequencing with mutant pexel-I/II of the *CSP* locus (1023 bp) was constructed by overlapping PCR and inserted into the multiple clone sites of the pYC plasmid. *P.b* ANKA CSP pexel domain I-II was mutated as previously reported.^1^ The amino acid sequence of pexel domain I, RNLNE, was changed to ANANA, while the sequence of pexel domain II, RLLAD, was changed to ALAAA. The resulting recombinant pYC-CSP_mut_ plasmid was amplified and purified using Endo-Free Plasmid Midi Kit (OMEGA, Norcross, GA, USA). Plasmid electro-transfection was performed as previously reported.^2^ In brief, Kunming mice were infected with *P.b* ANKA via intraperitoneal injection, and the erythrocytic-stage parasites were collected from the mice with a parasitemia of 5– 15%. The parasites were immediately added to a 250-ml conical flask containing 100 ml RPMI-1640 culture medium (HyClone, Logan, UT, USA) and 25 ml fetal bovine serum (FBS; Gibco, Detroit, MI, USA), and then a gas mixture of 5% CO2, 5% O2, and 90% N2 was flushed into the conical flask using a 0.22-μm filter (Millipore, Billerica, MA, USA). The conical flask was incubated at 37°C on a swing bed with shaking at a speed that was just high enough to keep the cells in suspension overnight. Subsequently, the schizonts were separated and collected by density gradient centrifugation with 72% Percoll (GE Healthcare, Amersham, Buckinghamshire, UK). Human T Cell Nucleofector^®^ Kit (Lonza, Basal, Switzerland) was used for electro-transfection. The schizonts were resuspended with 110 μL DNA solution including 100 μL Nucleofector solution and 10 μL ddH2O containing 10–20 μg recombinant pYC-CSP_mut_ plasmid and transfected using program U33. Fifty microliters of complete culture medium were added immediately and 150 μL of the complete transfection solution was injected into a tail vein of a Kunming mouse.

Once parasites appeared in the blood, the mouse was fed 8 μg/mL pyrimethamine (Sigma-Aldrich, St Louis, MO, USA) diluted in water to screen for mutant parasites. The pyrimethamine-resistant parasites were collected and cloned by intravenously injecting each mouse with 100 μL of a phosphate-buffered saline-diluted parasite solution containing ~1.0 infected iRBC. Nine to 12 days later, the blood of the mouse with cloned parasites was collected and the genomic DNA was extracted. The CSP pexel I-II mutation of the parasite clone was identified by PCR from the genomic DNA, followed by verification with DNA sequencing.

### Immunofluorescence assay

HepG2 cells (1 × 10^5^) were placed on a 14-mm-diameter slide in a 24-well plate. To detect the distribution of CSP, HepG2 cells were infected with WT or CSP_mut_ sporozoites. Three hours after invasion, the cells were washed with PBS to remove non-invaded sporozoites and incubated with fresh DMEM (Hyclone) containing 10% FBS (Gibco) and three antibiotics. Twenty-four hours after invasion, the slide was immobilized with 4% paraformaldehyde (Sangon Biotech) for 10 min, penetrated with 0.1% Triton-X 100 (Amersco, Albany, NY, USA) for 15 min, and then blocked with PBS containing 5% BSA, 0.02% Tween-20 for 1 h at room temperature. The cells were labeled with 1:500 goat anti-UIS4 (Sicgen, Cantanhede, Portugal) overnight at 4°C and 1:100 Green donkey anti-goat secondary antibody (Abbkine, Wuhan, China) for 1 h at room temperature. After washing with PBS three times, the cells were labeled with 1:500 rabbit anti-CSP overnight at 4°C and 1:100 Red donkey anti-rabbit secondary antibody (Abbkine) for 1 h at room temperature. The nuclei were counterstained with DAPI (Beyotime) for 5 min at room temperature.

For invasion rate observation, images were obtained at 6h after infection, UIS4 staining were used to recognize the parasites in HepG2 cells. For evaluating the co-localization of CSP_mut_ and LC3, the cells were transfected with hLC3B-RFP plasmid and infected with WT or CSP_mut_ sporozoites, and then labeled with goat anti-UIS4 and the secondary antibody as stated above. To observe the effect of CSP on the IFN-γ-mediated killing of EEFs, control and CSP-stably transfected HepG2 cells were pre-treated with 1 U/mL recombinant human IFN-γ (Peprotech, Rocky Hill, NJ, USA) or equivoluminal medium for 6 h. The cells were then infected with *P.b* ANKA or *P.b* ANKA-RFP as stated above and co-treated with IFN-γ 46 h hours after infection.

For examining effect of CSP overexpression on the co-localization of EEF and LC3, control and CSP-stably transfected HepG2 cells were transfected with hLC3B-RFP plasmid. Twenty-four hours after transfection, the cells were infected with *P.b* ANKA sporozoites as stated above for 24 h, and then treated as described above.

### Griess reaction analysis

The supernatant of the cell culture was collected 24 h after infection with sporozoites, transfected with pcDNA3.1 or pcDNA3.1-CSP, and treated with or without IFN-γ. For Griess analysis (Beyotime), the NaNO2 standard was diluted by DMEM containing 10% FBS to a final concentration of 0, 1, 2, 5, 10, 20, 40, 60, and 100 µM. Fifty microliters of different concentrations of NaNO2 standards and the supernatant were added to a 96-well plate, and then 50 μL of Griess Reagent I and II (Beyotime) were added in sequence at room temperature. After slight vibration, the concentration of NO was detected at 540 nm using iMark^TM^ Microplate Absorbance Reader (Bio-Rad). RAW264.7 cells stimulated with 100 ng/mL LPS (Sigma-Aldrich) were used as the positive control.

### Transcriptome sequencing

CSP-stably transfected or control cells (1 × 10^6^) were washed with PBS three times and lysed by 1 mL Trizol (Invitrogen). Total RNA was extracted as stated above. Qubit Fluorometric Quantitation 2.0 (Invitrogen) was used to detect the concentration of total RNA. A total amount of 2 μg RNA per sample was used as input material for the RNA sample preparations. Sequencing libraries were generated using VAHTSTM mRNA-seq V2 Library Prep Kit for Illumina^®^following the manufacturer’s recommendations, and index codes were added to attribute sequences to each sample. In brief, mRNA was purified from total RNA using poly-T oligo-attached magnetic beads. Fragmentation was carried out using divalent cations under elevated temperature in VAHTSTM First Strand Synthesis Reaction Buffer (5X). First-strand cDNA was synthesized using a random hexamer primer and M-MuLV Reverse Transcriptase (RNase H). Second-strand cDNA synthesis was subsequently performed using DNA polymerase I and RNase H. Remaining overhangs were converted into blunt ends via exonuclease/polymerase activities. After adenylation of the 3′ ends of DNA fragments, an adaptor was ligated for library preparation. To select cDNA fragments of preferentially 150–200 bp in length, the library fragments were purified with the AMPure XP system (Beckman Coulter, Beverly, USA). Then, 3 μL USER Enzyme (NEB, Ipswich, MA, UK) was incubated with size-selected, adaptor-ligated cDNA at 37°C for 15 min followed by 5 min at 95°C before PCR. PCR was performed with Phusion High-Fidelity DNA polymerase, Universal PCR primers, and Index (X) Primer. Finally, PCR products were purified (AMPure XP system) and the library quality was assessed on the Agilent Bioanalyzer 2100 system. The libraries were then quantified and pooled. Paired-end sequencing of the library was performed on HiSeq XTen sequencers (Illumina, San Diego, CA). RAW data have been submitted to GEO database, and series record is GSE129323.

### Total RNA extraction and real-time PCR

For parasite burden detection, the livers of mice were dissected at 46 h after infection of WT or CSP_mut_ sporozoites, and homogenized in 1.5 mL Trizol (Invitrogen), and HepG2 cells were collected 46 h after infection of WT or CSP_mut_ sporozoites and lysed by 1 mL Trizol (Invitrogen). 100μL liver homogenate or 1 mL of cell lysis were used for total RNA extraction using Trizol in accordance with the manufacturer’s instructions. cDNA was synthesized from equivalent total RNA using PrimeScript™ RT reagent Kit with gDNA Eraser (TAKARA) in accordance with the manufacturer’s instructions. The parasite load in the livers and HepG2 cells was evaluated by Taqman-PCR with primers and probes for 18S rRNA and *GAPDH* following the manufacturer’s instructions of Premix Ex Taq™ (Probe qPCR) (TAKARA). For the SYBR quantitative PCR assay, HepG2 cells in 6-well plates were lysed by 1 mL Trizol. Total RNA was extracted and cDNA was synthesized as described above. The mRNA levels of *IFN-γ*, *iNOS, LC3B, ATG3, ATG5, ATG7, STUB1, SYVN1, CBL, SMURF1, PAFAH1B1,* and *NEDD4* were evaluated using TB Green™ Premix Ex Taq™ II (Tli RNaseH Plus) (TAKARA). CFX96 Touch™ Real-Time PCR Detection System and CFX ManagerTM Software (Bio-Rad, Hercules, CA, USA) were used for RT-PCR data collection and analysis.

## Notes

**Funding** This work was supported by the National Natural Science Foundation of China (No. 81672053 and 81702247), the State Key Program of the National Natural Science Foundation of China (No. 81830067) and the Miaopu Talent Grant from Army Medical University (2019R057).

